# Cleavage modification did not alter early blastomere fates during bryozoan evolution

**DOI:** 10.1101/068783

**Authors:** Bruno C. Vellutini, José M. Martín-Durán, Andreas Hejnol

**Affiliations:** Sars International Centre for Marine Molecular Biology, University of Bergen, Thormøhlensgate 55, 5006 Bergen, Norway.

**Keywords:** Bryozoa, cyphonautes, spiral cleavage, cell lineage, larva, MAPK, gene expression, molecular patterning

## Abstract

Stereotypic cleavage patterns play a crucial role in cell fate determination by precisely positioning early embryonic blastomeres. Although misplaced cell divisions can alter blastomere fates and cause embryonic defects, cleavage patterns have changed several times during animal evolution. Here, we analyze the evolutionary transition from spiral cleavage – a stereotypic pattern remarkably conserved in many protostomes – to the biradial cleavage of bryozoans. We characterize the cell lineage, MAPK signaling and expression of several developmental genes in the bryozoan *Membranipora membranacea*, and found that the fate and the genes expressed in the early bryozoan blastomeres are similar to their putative homologous blastomeres in spiral-cleaving embryos. The data indicate that cleavage geometry evolved independent from other developmental traits during the transition from spiral to biradial cleavage in the bryozoan lineage, revealing that stereotypic cleavage patterns can be evolutionarily modified without major changes to the molecular identity and fate of embryonic blastomeres.

## Background

Cleavage is the sequence of cell divisions that turns a zygote into a multicellular embryo, and plays an essential role in the specification of cell fates before the onset of gastrulation. A cleavage pattern can be variable, where the blastomere positions are not predictable (e.g. mouse), or stereotypic (e.g. ascidian), where the embryonic cell divisions form a precise, identifiable three-dimensional pattern of blastomeres (Davidson, 1990). There is evidence that different types of cleavage can dictate different underlying mechanisms of cell fate specification (Davidson, 1990), however, it is still largely unknown how a stereotypic cleavage pattern affects the evolution of animal morphology (Wray, 1994). Cleavage patterns are highly diverse, they can even differ between closely related species (Raff, 1992; Schulze and Schierenberg, 2011), or remain conserved in different animal lineages over long evolutionary periods (Hejnol, 2010). A notable example of the latter is known as *spiral cleavage*, and is a rich framework to investigate the relation between development and evolution.

Spiral cleavage occurs in molluscs, annelids, nemerteans and polyclad flatworms (Child, 1897; Child, 1900; Conklin, 1897; Eisig, 1898; Lillie, 1895; Nelson, 1904; Surface, 1907; Treadwell, 1901; Wilson, 1892; Wilson, 1903; Zeleny, 1904). In these groups, the fertilized eggs divide through a highly stereotypic cleavage pattern where blastomeres at the 4-cell stage cleave with the mitotic spindles oblique to the animal-vegetal axis, alternating direction (clockwise and counterclockwise) at each division cycle – the spiral cleavage pattern (Costello and Henley, 1976; Hejnol, 2010; Henry and Martindale, 1999; Lambert, 2010; Wilson, 1892). This determinate developmental mode allowed for the identification of homologous blastomeres across taxa and unprecedented detail in the comparison of animal embryogenesis, further revealing that spiral-cleaving embryos not only have the same cleavage pattern, but homologous blastomeres between groups have a similar fate in the larval and adult tissues (Hejnol, 2010; Henry and Martindale, 1999). The study of spiral cleavage thus revealed that early development can remain conserved for extended evolutionary periods, in contrast to late developmental stages, shaping our current understanding about the relation between ontogeny and phylogeny (Dohle, 1989a; Dohle, 1989b; Guralnick, 2002; Maienschein, 1978; Scholtz, 2005).

Even though spiral cleavage has been modified in a multitude of ways throughout evolution, with changes in blastomere sizes and cell fate specification (Costello and Henley, 1976; Hejnol, 2010; Henry and Martindale, 1999; Lambert, 2010), the cleavage pattern itself remained fairly conserved. Known cases where the spiral cleavage pattern was lost is usually associated to drastic developmental changes, such as the transition to a syncytial blastoderm in cephalopods (Wadeson and Crawford, 2003), or to the evolution of extra-embryonic yolk cells in platyhelminthes (Martín-Durán and Egger, 2012). However, the recent improvements in the resolution of protostome relationships reveal that the spiral cleavage pattern has been drastically modified or lost in even more groups than previously thought (Hejnol, 2010).

Phylogenetic analyses suggest that spiral-cleaving groups belong to the Spiralia, a major protostome clade sister to the Ecdysozoa (e.g. insects and nematodes) (Dunn et al., 2014). Some spiralians (i.e. animals belonging to the clade Spiralia) such as gnathostomulids (Riedl, 1969), phoronids (Pennerstorfer and Scholtz, 2012) and entoprocts (Marcus, 1939; Merkel et al., 2012), also display a spiral-like cleavage. Other clades that have been recognized as spiralians, such as bryozoans (Santagata, 2015a), brachiopods (Santagata, 2015b), gastrotrichs (Hejnol, 2015a) and rotifers (Hejnol, 2015b), do not show spiral cleavage (Figure 1). Mapping the spiral geometry to recent spiralian phylogenies (Dunn et al., 2014; Kocot, 2016; Kocot et al., 2016; Laumer et al., 2015; Nesnidal et al., 2013; Struck et al., 2014) indicates that spiral cleavage is a synapomorphy for the Spiralia, or at least for the Lophotrochozoa+Rouphozoa clade (Figure 1). This implies that the spiral cleavage pattern was modified during the evolution of gastrotrichs, brachiopods and bryozoans (Dunn et al., 2008; Hejnol, 2010). Therefore, these groups are essential to understand how cleavage patterns and blastomere fates evolve and can uniquely reveal which developmental traits, if any, remained conserved in the evolutionary transition from spiral to a derived cleavage geometry.

**Figure 1.**
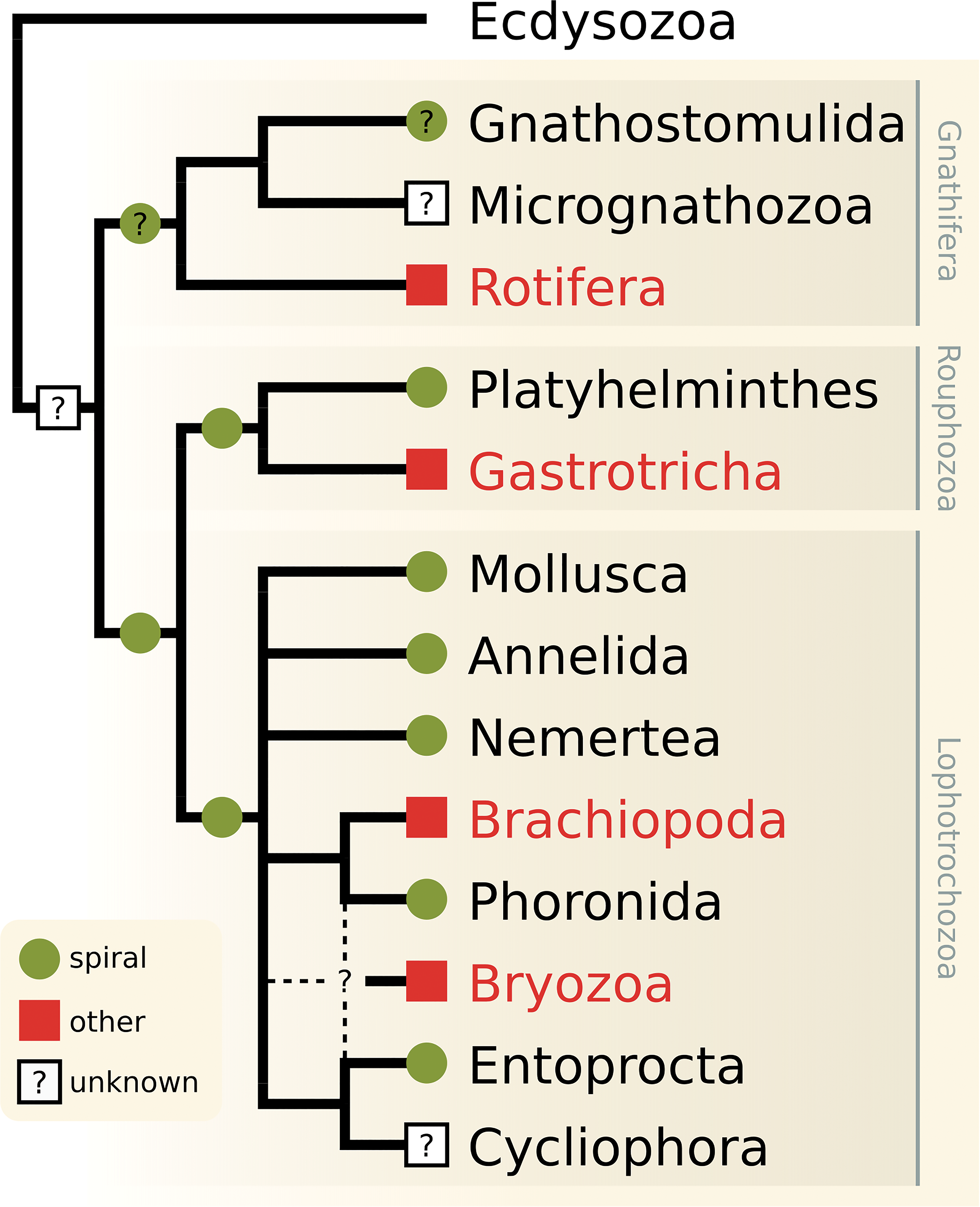
Phylogenetic distribution of the spiral cleavage pattern in the Spiralia. Symbols indicate taxa containing spiral-cleaving species (green circles), taxa where the spiral cleavage geometry has not been identified (red squares) and taxa where the cleavage pattern is unknown (white square with question mark). Symbols placed on nodes mark the presumed ancestral cleavage pattern of the branch. Green circles with question mark indicate that the presence of a spiral cleavage pattern is uncertain (i.e. Gnathostomulida and Gnathifera). Spiralian relationships based on Nesnidal et al. (2013), Struck et al. (2014) Laumer et al. (2015), Kocot et al. (2016) and Kocot (2016), and cleavage data based on Hejnol (2010). Dashed lines indicate alternative placements for Bryozoa.

In the current work, we investigate the development of a group that lost the spiral cleavage pattern during evolution – the bryozoans. These sessile colonial invertebrates occur in oceans worldwide and have fairly distinct developmental patterns regarding reproduction (e.g. brooding versus external development), early development (e.g. cleavage and gastrulation) and larval stages (Reed, 1991; Zimmer, 1997), but none of the species investigated so far display a spiral arrangement of embryonic blastomeres (Reed, 1991). Yet, bryozoans display a unique stereotypic cleavage pattern with a biradial arrangement of the blastomeres that is widely conserved within the group.

Early embryological studies (Barrois, 1877; Calvet, 1900; Corrêa, 1948; Marcus, 1938; Pace, 1906; Prouho, 1892) suggest that the animal-most blastomeres of gymnolaemate embryos originate the apical disc and aboral epithelium of the larva, the animal micromeres at the equator of the embryo form the ciliated band, and the vegetal cells produce the oral epithelium and endomesoderm (Hyman, 1959; Reed, 1991; Zimmer, 1997). A cell lineage topology deemed comparable to the cell lineage of spiral-cleaving embryos (Nielsen, 2001; Nielsen, 2005). However, cleavage patterns have only been systematically followed until the 64-cell stage (Corrêa, 1948; Pace, 1906) and, as of today, there is no detailed cell lineage of a bryozoan larva. Several basic developmental questions remain unsolved. For example, the relation between the embryonic animal/vegetal axis and the larval body axes is unclear (Nielsen, 2005), and the fate of the blastopore remains to be confirmed (Gruhl, 2009; Marcus, 1938; Prouho, 1892; Zimmer, 1997). Finally, the fate of internalized cells has not been traced (Zimmer, 1997) and the source of mesoderm remains a specially contentious topic (Gruhl, 2009).

In this study, we investigate the embryogenesis of the cosmopolitan gymnolaemate species *Membranipora membranacea* (Linnaeus, 1767) to understand the evolutionary transition from a spiral to a biradial cleavage pattern. We take advantage of the vast cell lineage data available for spiralians and the growing literature on spiralian gene expression, to compare with cellular resolution the molecular identity and fate of embryonic blastomeres between the bryozoan and other spiral-cleaving embryos. We were able to identify the embryonic source of most larval tissues of *M. membranacea* based on 4D microscopy recordings, and to combine this cell lineage data with the activity of the MAPK pathway and expression of several conserved developmental markers, generating a detailed overview of the blastomere identities and fates in the bryozoan. The comparison to a typical spiral development reveals that the early blastomeres of *M. membranacea* share similar molecular identities and fates with other spiral-cleaving embryos, despite the contrasting cleavage pattern. Given the phylogenetic position of bryozoans, we suggest these coincident developmental traits were inherited from a spiral-cleaving ancestor during the evolutionary transition from spiral to biradial cleavage. The findings support the hypothesis that stereotypic cleavage patterns can be modified during evolution without major changes to blastomere gene expression and fates. Our study highlights the power of the comparative approach to address fundamental questions of development and evolution, such as the relation between cleavage patterns and fate maps.

## Results

### General development

Colonies of *M. membranacea* spawn fertilized discoidal eggs into the water column (Temkin, 1994). The released eggs undergo activation, becoming spherical (Figure 2A), and initiate cleavage with a discernible accumulation of yolk in the vegetal pole (Figure 2B). Throughout development, the embryo maintains close contact with the fertilization envelope via abundant cytoplasmic extensions (Figure 2A; Figure 2I). The yolky cells at the vegetal pole are internalized during gastrulation (Figure 2C–E; Figure 2J–M) and, by mid-gastrula stage, the primordium of the apical organ (apical disc) and of the ciliated band (corona) are visible (Figure 2E). The vegetal plate invaginates and the embryo elongates along the animal-vegetal axis forming a late gastrula with clearly defined larval structures (i.e. apical organ, shell, gut and corona) (Figure 2F–G). At this point the fertilization envelope opens at the apical and basal ends and the embryo begins to swim by ciliary beating (Figure 2F). The internal cavity (vestibule) widens in the anteroposterior axis resulting in the typical laterally compressed, triangular shaped and shelled feeding larva of gymnolaemate bryozoans – the cyphonautes (Figure 2H–I) (Stricker et al., 1988). In this study, we recorded and traced the cell lineage from the 2-cell stage (Figure 2B) until the mid/late gastrula stage (Figure 2E–G).

**Figure 2.**
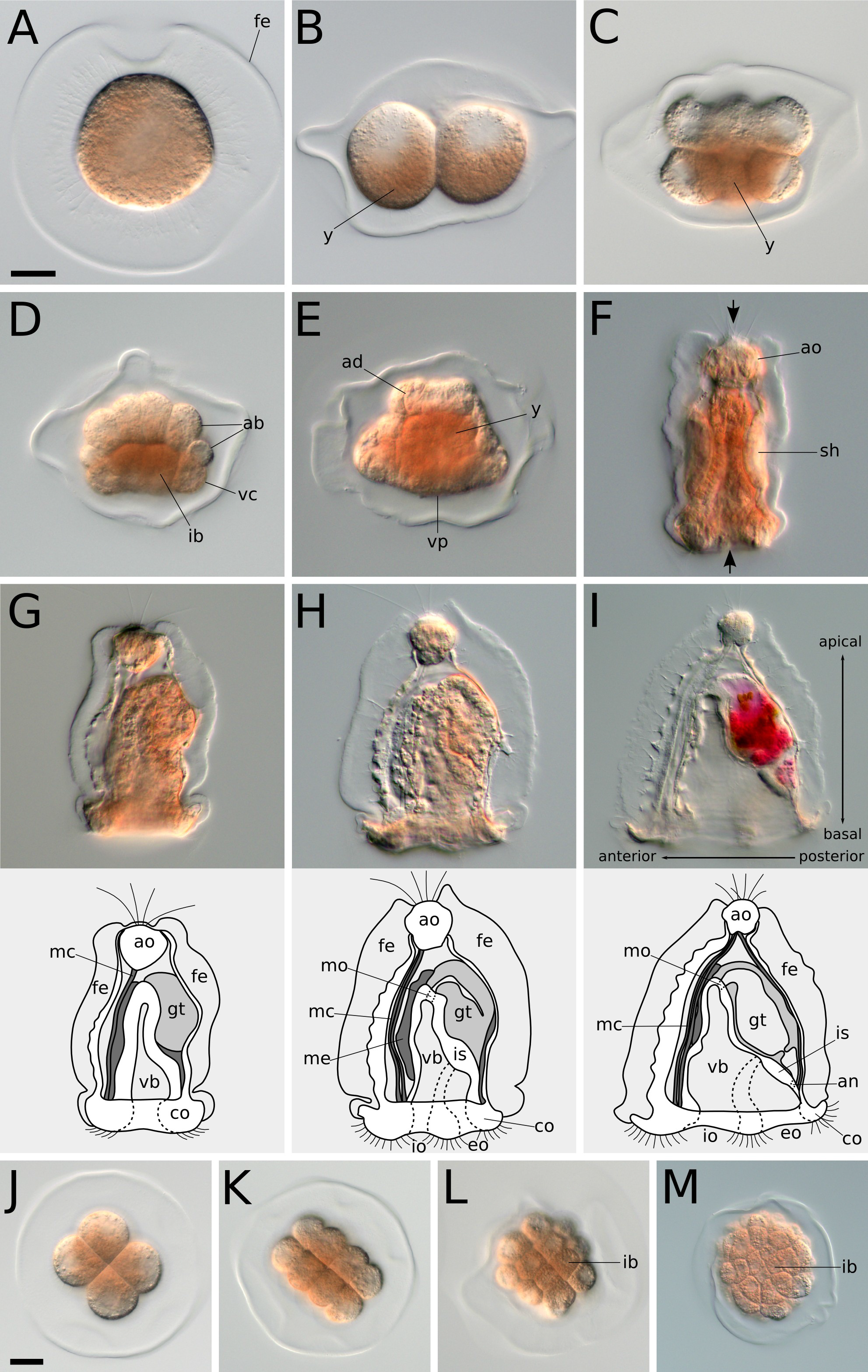
Overview of *M. membranacea* development. (A) Vegetal view of an activated egg becoming spherical. (B–I) Animal pole is top and vegetal pole is bottom. (B) 2-cell stage showing higher amount of yolk (y) on the vegetal side. (C) 8-cell stage with yolk positioned on the inner cytoplasmic portions. (D) 28-cell stage. Large vegetal blastomeres carry most of the yolk with less yolk in the animal blastomeres (ab). (E) Anterior view of a mid-gastrula stage with a prominent apical disc (ad), vegetal ectoderm (ve) and an inner mass of yolk-rich blastomeres. (F) Frontal view of a late gastrula stage after the vegetal ectoderm invaginated and the embryo extended in the animal/vegetal axis. Most larval structures are formed by this stage, including the apical organ (ao) and shell valves (sh). At this stage, the cilia of the apical tuft and the corona breakthrough the fertilization envelope (arrows). (G–I) Lateral views showing the morphogenesis of the cyphonautes for 24h late gastrula (G), 48h larva (H) and 3d larva (I). Larval structures are illustrated below each panel. (J–M) Vegetal views showing beginning of gastrulation. (J) 8-cell stage. (K) 16-cell stage. (L) 28-cell stage. (M) 90-cell stage with vegetal blastomeres being internalized. co: corona, gt: gut, vb: vestibule, mc: muscle cell, me: mesodermal tissue, is: internal sac, mo: mouth, io: inhalant opening, eo: exhalant opening, an: anus. Scale bars = 20 µm.

### Cleavage pattern and embryonic axes

The cleavage of *M. membranacea* is biradial as previously described for gymnolaemates (Figure 3) (Gruhl, 2009; Nielsen, 2005; Reed, 1987; Reed, 1991; Zimmer, 1997). At 15 °C, the first cell division occurs between 1–2 hours post activation (hpa) and originates two equal blastomeres with a meridional cleavage furrow. The second division is also meridional and perpendicular to the first, resulting in four identical blastomeres at 3 hpa with an unequal animal/vegetal distribution of yolk (Figure 2B; Figure 3; Figure 4, 4-cell). We labeled the blastomere that originates the posterior structures of the larval body as “D” (see Figure 5 for fate map overview and “Methods” for nomenclature details). In most embryos, the cell sister of the D blastomere gives rise to the right side of the embryo (Martín-Durán et al., 2016a). At 4 hpa, an equatorial third division gives rise to four animal blastomeres with little yolk content (1a–1d) and four equally sized vegetal blastomeres with a greater amount of yolk in the central portion of the embryo (1A–1D) (Figure 2C; Figure 3; Figure 4, 8-cell). On the next division at 5.2 hpa, each animal blastomere divides parallel to the plane of the first cleavage resulting in four internal (subscript i) and four external (subscript e) cells; vegetal blastomeres cleave in the same manner slightly after (Figure 3; Figure 4, 16-cell). At the 16-cell stage, yolk-rich cells (2A–2D) lie inner to the outer vegetal cells of the second quartet (2a–2d) and the embryo is clearly biradial.

**Figure 3.**
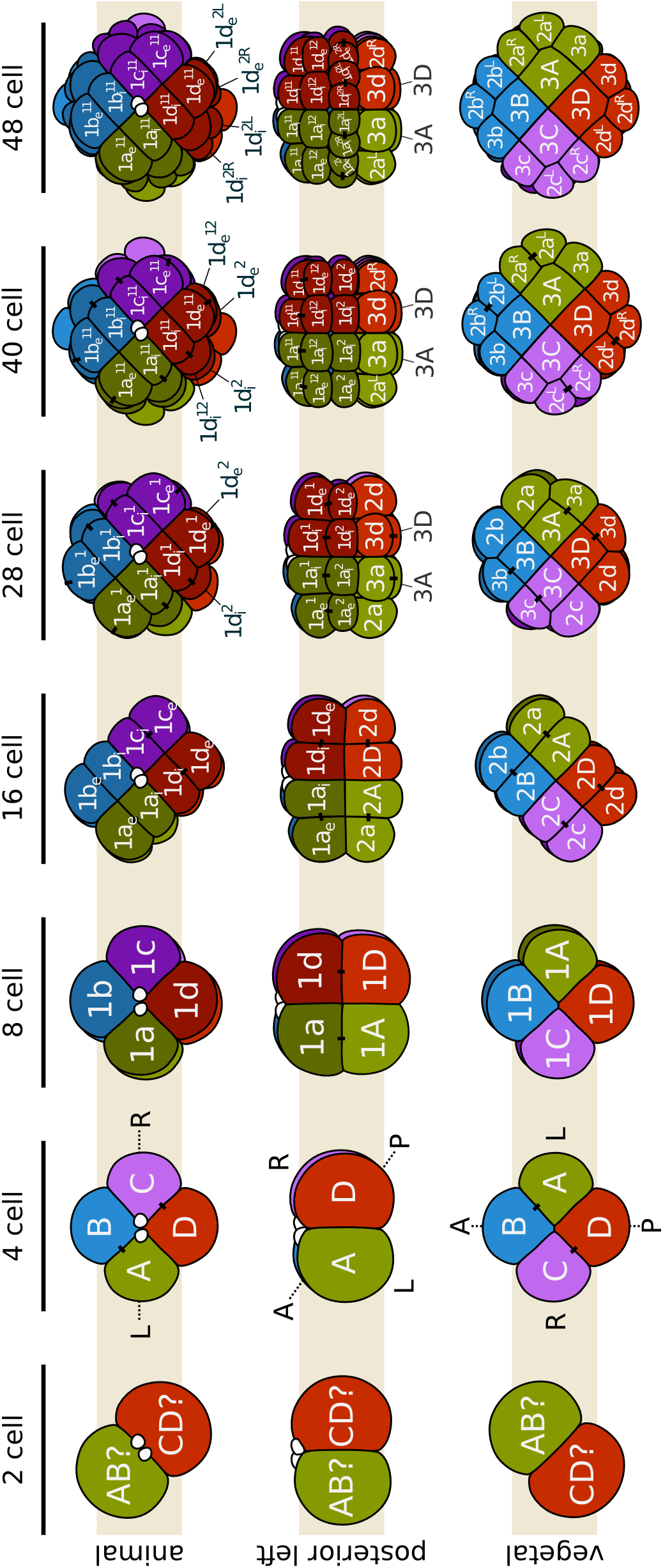
Cleavage pattern and orientation of the embryonic axes of *M. membranacea*. Quadrant identity was determined backwards from 4D microscopy recordings and it is not known if it is determined during early stages.

**Figure 4.**
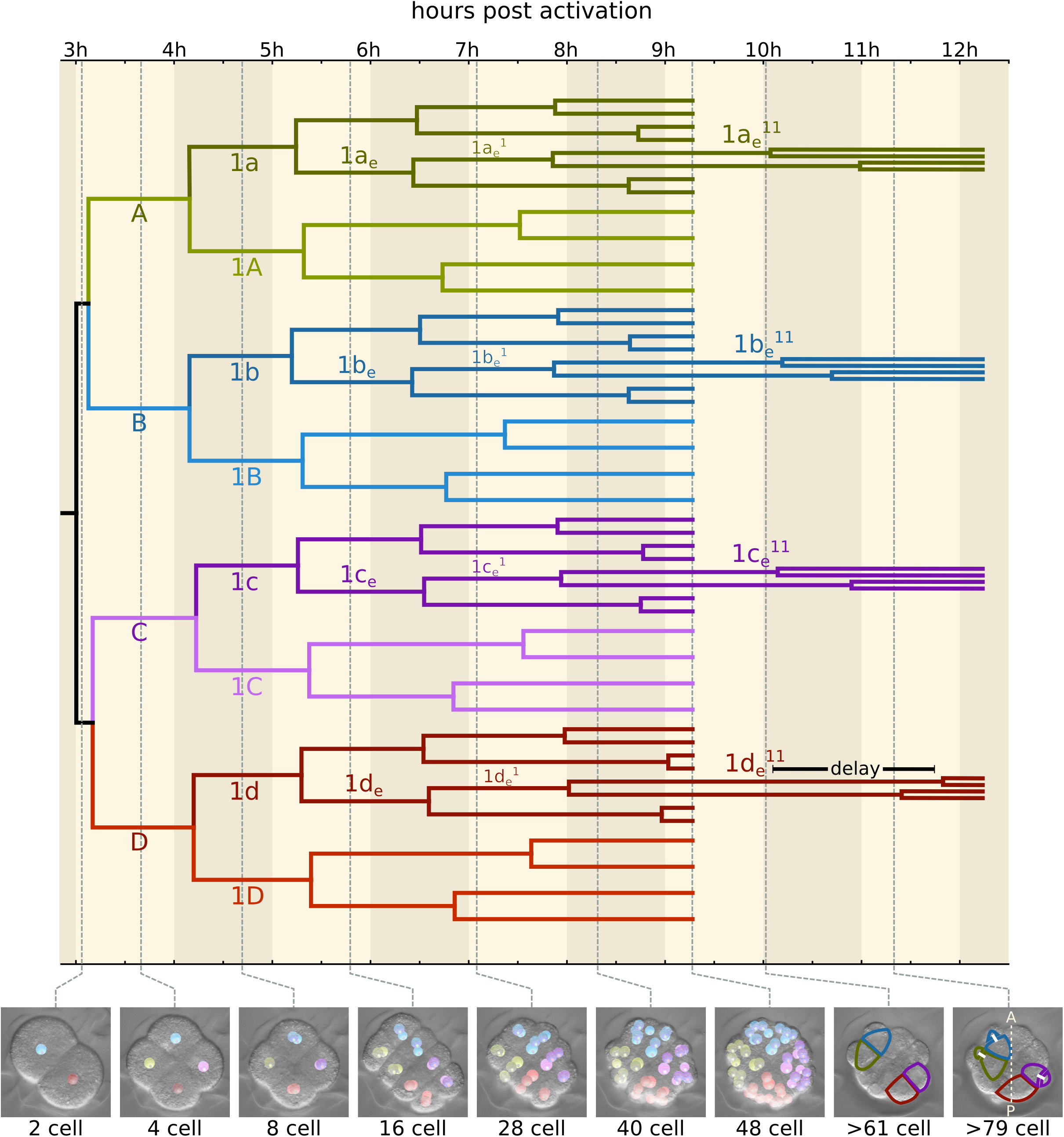
Timed cell lineage of *M. membranacea* and the break of biradial symmetry. Panels onthe bottom show the developmental stages with the cell tracing overlay until the embryo with 48 cells. The outlines in the last two panels (>61 and >79 cells) indicate the cells 1a_e_^11^–1d_e_^11^, and the prominent delay in the division of 1d_e_^11^. The anteroposterior axis is denoted by a dashed line in the last panel. Quadrant color coding: A (green), B (blue), C (purple), D (red).

**Figure 5.**
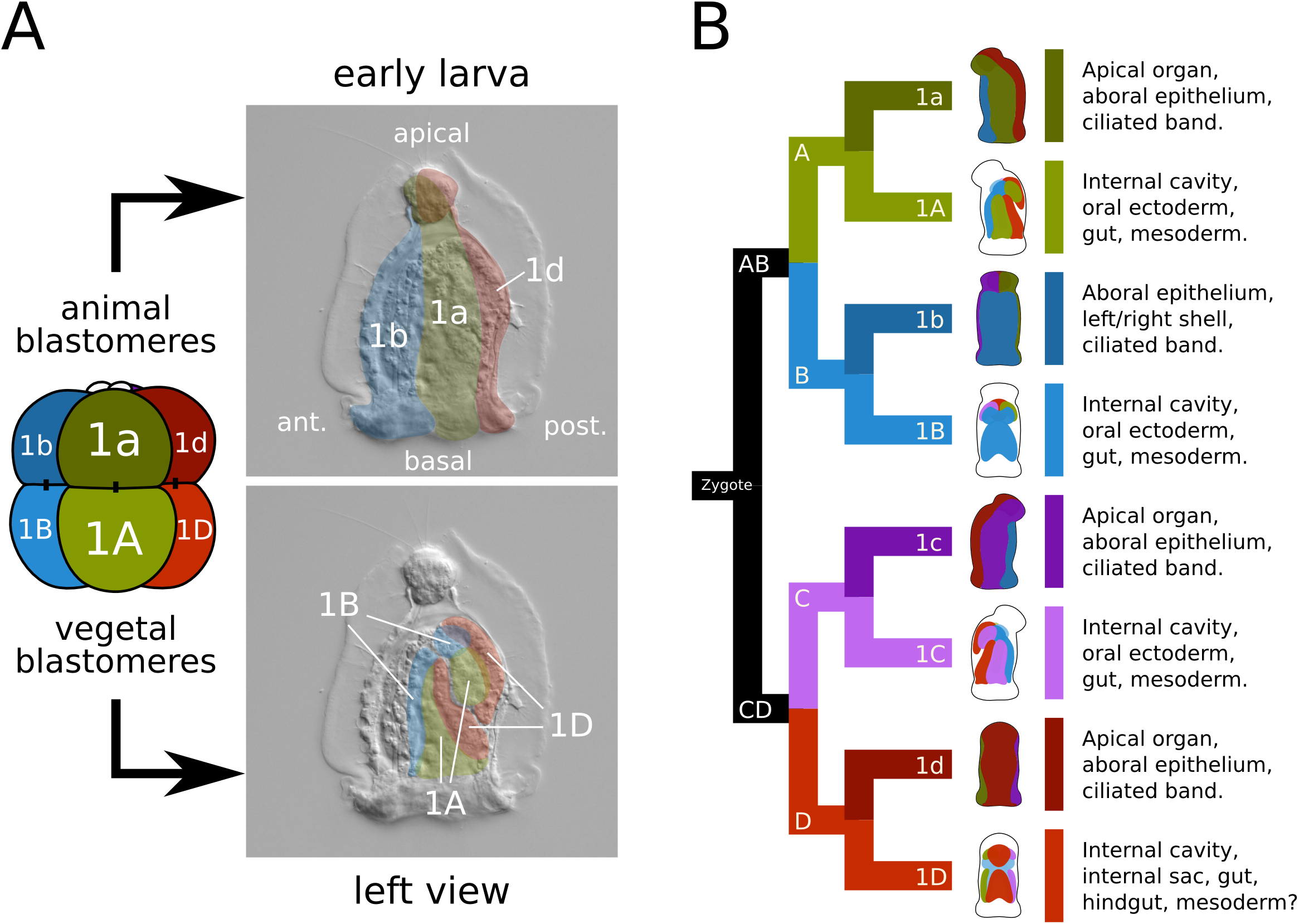
Larval fates of the eight embryonic blastomeres of *M. membranacea*. (A) Illustration based on the cell lineage data representing the overall fates of the animal (1a–1d) and vegetal (1A–1D) blastomeres. Animal blastomeres give rise to the apical organ, aboral epithelium and corona. Vegetal blastomeres give rise to the vestibule epithelium, oral ectoderm, mesoderm and gut. (B) Larval structures derived from each of the eight blastomeres. Quadrant color coding: A (green), B (blue), C (purple), D (red).

On subsequent stages, the eight animal blastomeres of *M. membranacea* act as octets, synchronized in the cell divisions (Figure 3). The first octet (animal pole cells 1q, 4 inner and four outer cells) divides equatorially making a brief 24-cell stage and the octets 1q^1^ and 1q^2^ (6.5 hpa). This division is shortly followed by an unequal cleavage originating the third quartet (3a–3d) from the four inner vegetal blastomeres at 6.8 hpa (Figure 3; Figure 4, 28-cell). Outer vegetal cells of the second quartet (2q) divide parallel to the second division at 7.5 hpa, resulting in twelve outer vegetal cells (3a–3d, 2a^R^–2d^R^, 2a^L^–2d^L^) that surround four large blastomeres in the vegetal plate (3A–3D) at the 32-cell stage. At 8 hpa, the top animal octet (1q^1^) divides forming a 40-cell embryo (Figure 3; Figure 4, 40-cell). Finally, the vegetal most animal octet (1q^2^) divides meridionally at 8.6–9 hpa forming an equatorial row of cells above the vegetal blastomeres (Figure 3; Figure 4, 48-cell).

Cell divisions between the correspondent blastomeres of each quartet are mostly synchronous up to the 64-cell stage (Figure 4). At this point, we observe the fist two significant asynchronous cell divisions in the posterior D quadrant (Figure S1). The cell 1d_i_^12^ divides approximately 1h before than its partners 1a_i_^12^, 1b_i_^12^ and 1c_i_^12^ (Figure S1), while the cell 1d_e_^11^ divides 2h later than its quartet correspondents 1a_e_^11^, 1b_e_^11^ and 1c_e_^11^ (Figure 4 and Figure S1). Other cells from the D quadrant also exhibit asynchrony, such as the large inner vegetal blastomeres 3Q (3D is delayed) and the outer vegetal cells 3q (3c is delayed). At a similar time point we observe the first difference in the orientation of the cleavage plane between quartet cells, whereas 1d_e_^12^ divides equatorially, 1a_e_^12^, 1b_e_^12^ and 1c_e_^12^ divide meridionally. These events are the first morphological evidence for the break in the biradial symmetry of the bryozoan embryo.

### Cellular origin of larval tissues

The larval body of *M. membranacea* develops from the four quadrants in a symmetrical manner, each lineage contributing almost equally to the structures on their respective sides: D=posterior, C=right, B=anterior and A=left (Figure 5A).

Progeny of the first quartet of animal blastomeres (1a–1d) originates apical ectodermal structures such as the apical organ, the aboral epithelia and the corona (Figure 5A). The apical organ is derived from derivatives of the apical-most cells 1a^1^, 1c^1^ and 1d^1^ (Figure 5B; Figure 6A). Cells 1a and 1c form the lateral and anterior most portion of the apical organ while the posterior cell 1d contributes not only to the posterior portion, but also to the tissues at the base of the apical organ (Figure 6A). Thus, the cell 1b is the only blastomere of the first animal quartet that does not contribute to the apical organ. Epithelial cells between the apical organ and the corona are mostly derived from the octets 1q^11^ and 1q^12^. Outer coronal cells originate from 1q^12^ and 1q^2^ while inner coronal cells (turned inwards after the invagination of the vegetal plate) are derived from 1q^2^ (Figure 6A). For a detailed overview of cell fates see Figure S2.

**Figure 6.**
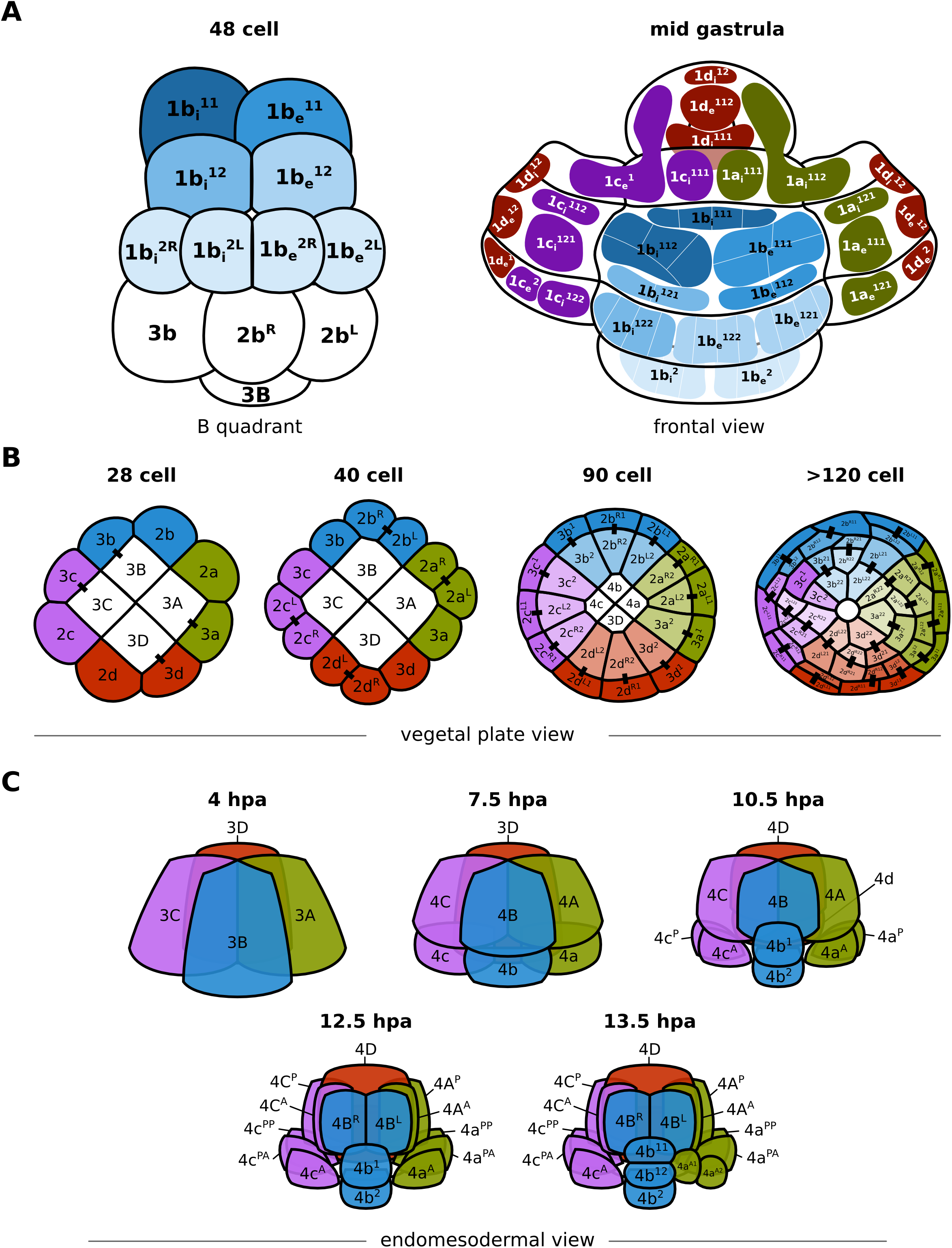
Details of the fate map and cleavage patterns of the animal blastomeres, vegetal ectoderm and internalized blastomeres of *M. membranacea*. (A) Representation of the B quadrant in frontal view at the 48-cell stage. Frontal view of the embryo at the mid gastrula stage with left, right and top regions “opened” for visualization. Shades of blue on the blastomeres of the 48-cell stage correspond to the fates in the epithelium and corona. White lines illustrate cell borders of further progeny from the blastomeres indicated. The fates of the blastomeres of other quadrants are indicated with the standard colors. (B) Cleavage patterns at the vegetal ectoderm. The original twelve-tet at the 40-cell stage divides once forming twelve derivatives, lining the forming blastopore at the 90-cell stage. At the subsequent division (>120 cells), progeny from the cells at the vertices (2a^L2^, 2b^R2^, 2d^R2^ and 2c^L2^) disconnect from the blastopore lip. At this stage, only eight cells are lining the blastopore. The cell 3c^1^ and 3c^2^ do not divide. (C) Cleavage patterns of the four large blastomeres that internalize during gastrulation. Quadrant color coding: A (green), B (blue), C (purple), D (red).

The vegetal blastomeres 1A–1D form the epithelium of the vestibule, the oral/anal ectoderm as well as the cells internalized during gastrulation, which originate the endoderm and mesoderm of the cyphonautes larva (Figure 5A). The cellular arrangement at the vegetal plate in a 40-cell embryo consists of 12 outer cells (3a–3d, 2a^R/L^–2d^R/L^) and four large inner blastomeres (3A–3D) (Figure 6B). Here we define gastrulation as the internalization of these four vegetal cells. It occurs by delamination and epiboly in two rounds of division of the outer vegetal twelve-tets, which divide radially, pushing the 4 larger blastomeres internally and outlining a blastopore (Figure 6B, 90 cell). At the 90-cell stage 12 cells are defining the blastopore lip, but this number gets reduced to 8 cells after the next division (Figure 6B, 120 cell). From the 12 vegetal cells, one does not divide (3c^2^) and continue to line the right side of the blastopore lip (Figure 6B, 120 cells). Cells at the vertices of the blastopore on the 90-cell stage (2a^L2^, 2b^R2^, 2c^L2^ and 2d^R2^) get pushed away from the blastopore lip, which now consists of 8 cells (Figure 6B). Blastomeres not forming the blastoporal lip also undergo the same round of two radial divisions except for 3c^1^, the sister of 3c^2^. The derivatives of these 12 vegetal outer blastomeres form the whole ectoderm that invaginates and develops into the epithelia of the vestibule and preoral funnel. Thus, during the invagination of the vegetal plate and the apical-basal elongation of the embryo, the fate of the blastopore in *M. membranacea* is the larval mouth.

During epiboly (embryo around 90 cells), three of the internalized large blastomeres (3A–3C) undergo a round of unequal division forming the basal cells 4a–4c and the apical cells 1A–1C (Figure 6C). The division of the cell 3D occurs 3h delayed in comparison to the other blastomeres. This round of division sets apart the endoderm (4A–4D) from the mesodermal tissues (4a–4d) of *M. membranacea* cyphonautes larva. The cells 4a and 4c divide twice anteroposteriorly forming a pair of lateral rows of mesodermal cells (Figure 6C). The most anterior cells (4a^A^ and 4c^A^) form a bilateral pair of muscle cells extending from the corona to the apical organ (Figure 7). At the frontal portion of the larva, the cell 4b divides forming a column of cells stacking from the corona until the apical organ; the identity or role of these cells is unknown (Figure 6C). We could not resolve the fate of the 4d cell. Blastomeres 4A and 4C undergo anteroposterior divisions while 4B divides meridionally lining up with the blastoporal opening and forming the endodermal tissues of the cyphonautes larva (Figure 6C; Figure 7).

**Figure 7.**
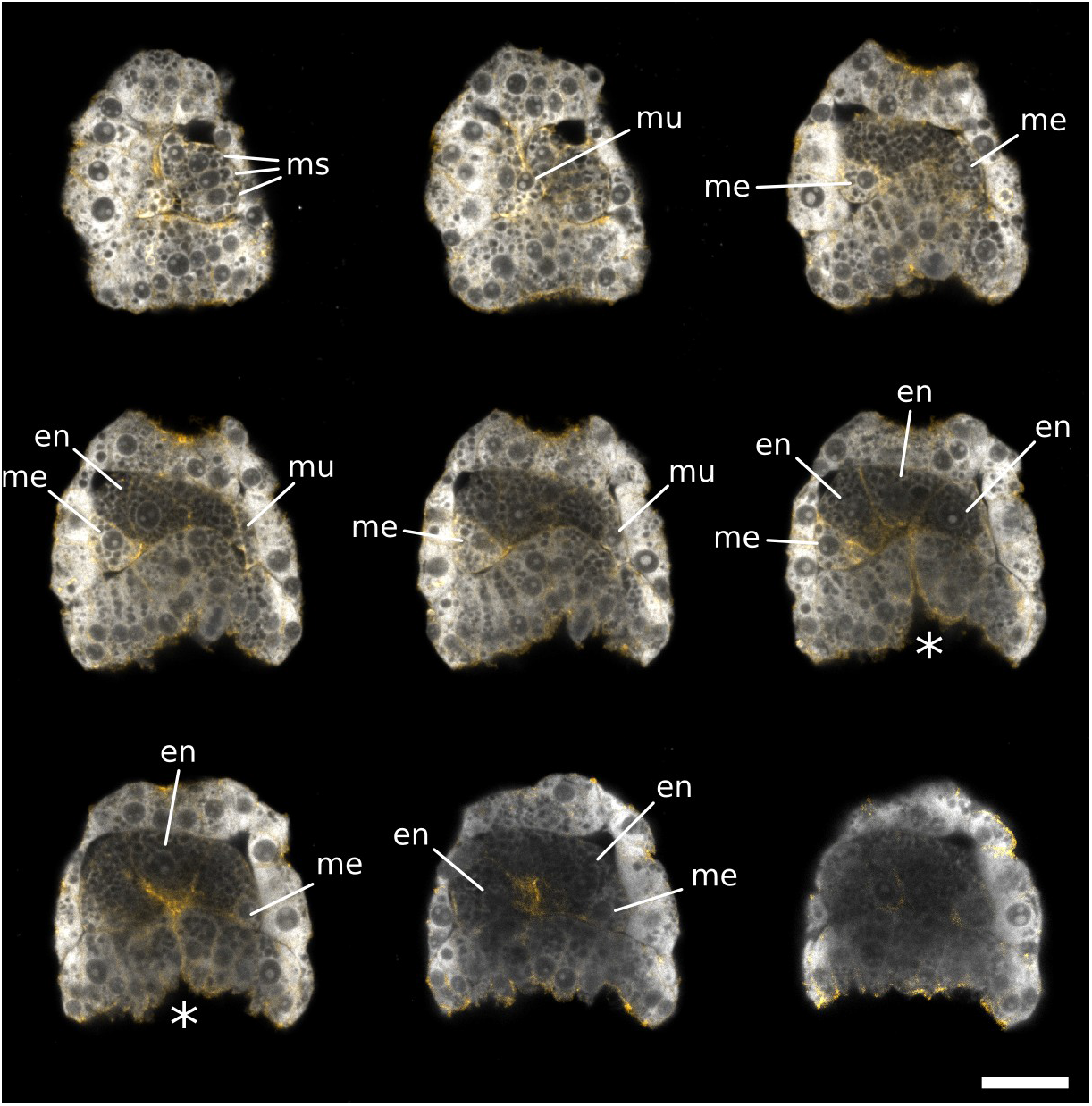
Mesodermal and endodermal cells in *M. membranacea*. Maximum intensity projections of 2–3 slices from a confocal stack at mid-gastrula stage. View of the anterior/right side of the embryo (top-left) to the posterior/left side (bottom-right). Samples stained with propidium iodide (grays) and phallacidin (orange). ms: mesodermal stack of cells from the B quadrant (4b derivatives), mu: muscle cells reaching the apical organ, me: other mesodermal cells, en: endodermal cells. Asterisk indicates the blastopore. Scale bar = 20 µm.

### MAPK activity

Previous work revealed that the MAPK signaling pathway might establish the position of the dorsal organizer in molluscan embryos (Lambert and Nagy, 2001). So far, all investigated molluscs show the asymmetric activation of MAPK in the 3D blastomere (Henry and Perry, 2007; Koop et al., 2007; Lambert and Nagy, 2001; Lambert and Nagy, 2003). Using an antibody against the activated form of MAPK, we found that in the bryozoan *M. membranacea* the first detectable MAPK activity also occurs in the 3D vegetal blastomere at the 28-cell stage (Figure 8B). MAPK activity persists in the 3D cell from the 28–90-cell stage and fades prior to the 3D division around 90-cell stage (Figure 8B–D). MAPK activity is not continued in the progeny of 3D, 4D or 4d (Figure 8E–F) and was not detected in later embryonic stages.

**Figure 8.**
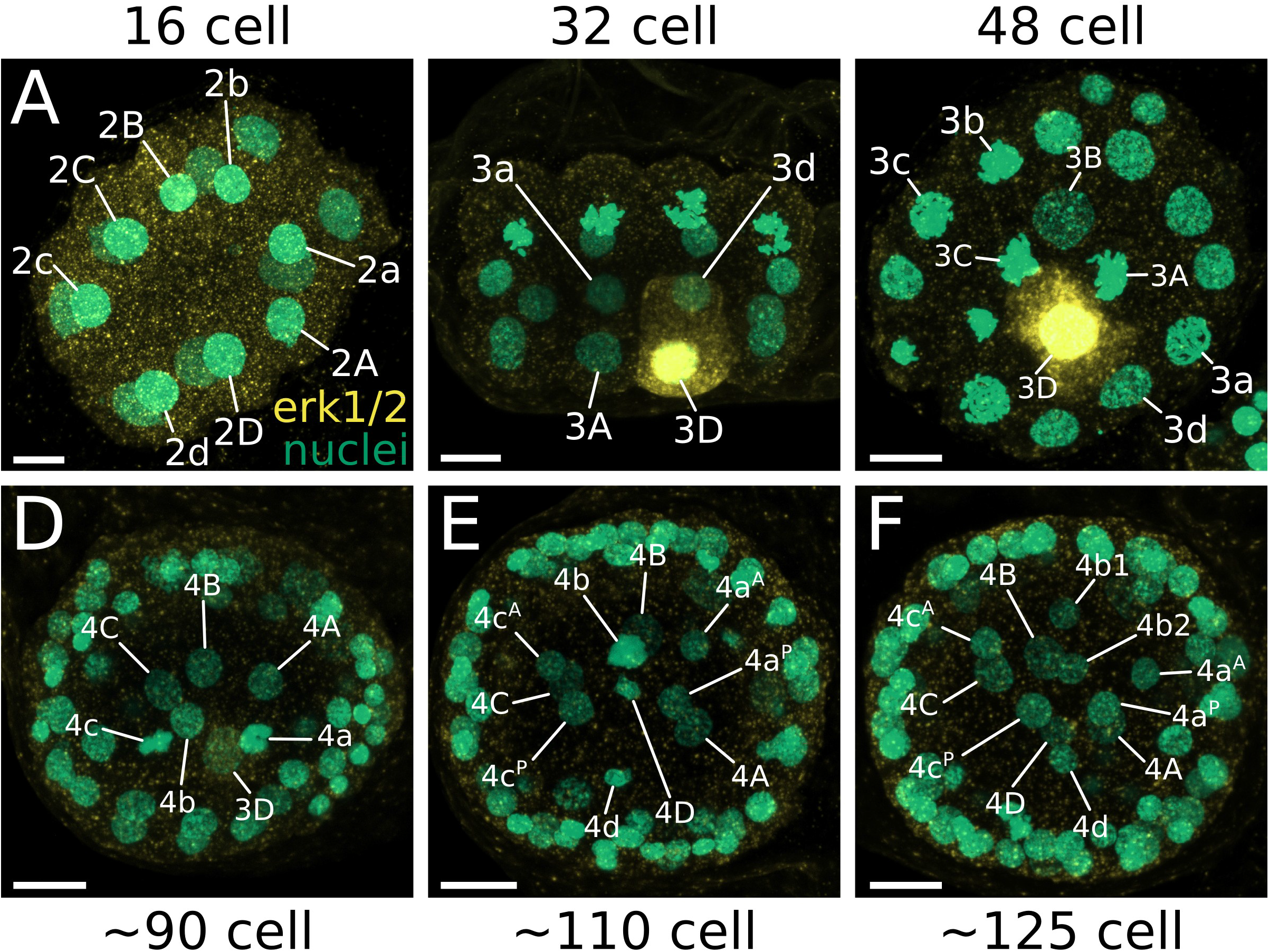
MAPK activity during the development of *M. membranacea*. (A) No detectable levels of activated MAPK at the 16-cell stage. (B) Side view showing only the quadrants A and D of a 32-cell stage with activated MAPK in the cell 3D. (C) Vegetal view of a 48-cell stage with blastomeres 3C and 3A undergoing mitosis. (D) Frontal view of an embryo with approximately 90 cells. 3D cell shows a weaker signal for MAPK activity. (E) Embryo soon after the division of the 3D cell. There are no detectable levels of MAPK activity in any cell. (F) Embryo with more than 125 cells without any detectable levels of MAPK activity. Scale bar = 10 µm.

### MAPK inhibition

The inhibition of the MAPK pathway in molluscs causes defects in the dorsoventral patterning (Henry and Perry, 2007; Koop et al., 2007; Lambert and Nagy, 2001; Lambert and Nagy, 2003), while in annelids MAPK-inhibited embryos have disorganized muscle and nerve tracts and overall shortened morphology (Amiel et al., 2013; Kozin et al., 2016; Pfeifer et al., 2014). We used the MEK inhibitor U0126 to investigate the role of the MAPK pathway in the development of *M. membranacea*.

We investigated the effects of different U0126 concentrations (1, 10, 25 µM) on the development of *M. membranacea* when applied at the 2-cell stage (Figure S3A). We found the severity of the phenotype correlates with the concentration of the inhibitor, where the higher concentrations of 10 and 25 µM result in the complete disruption of the normal morphology (Figure S3A). These embryos show no identifiable larval structures, such as a differentiated apical organ or musculature, have a lower number of cells, and are shorter compared to control samples (Figure S3B). The percentage of this severe phenotype decreases when we begin the treatment at later developmental stages (from 4–8 hpa), and embryos treated from 10 hpa onwards show progressively milder phenotypes (Figure S4). In treatments beginning at 10–16 hpa, the larval structures such as apical organ, ciliated band and gut, are formed but the embryos are shorter and delayed in development in comparison to control embryos, while 18–24 hpa samples have almost normal morphology (Figure S4). Finally, to identify the developmental defect caused by the MEK inhibitor in the severe phenotype, we recorded *M. membranacea* embryos treated with 10 µM U0126 under the 4D microscope. We found that embryos with milder phenotypes develop slower when compared to wild type, but do not show any obvious cleavage abnormalities (Figure S5). However, embryos exhibiting the severe phenotype have a misguided 4th cleavage (8–16 cell stage), where the animal blastomeres no longer divide in a biradial fashion (Figure S5). Thus, the developmental defect causing the severe phenotype in U0126-treated embryos occurs before the 28-cell stage, when we first detect MAPK activity in *M. membranacea* (Figure 8).

### Gene expression

In order to complement the cell lineage studies and elucidate the blastomere identities of the bryozoan embryo, we analyzed the expression of sixteen conserved molecular markers during the development of *M. membranacea* (Figure 9–11; Figure S6): anterior/neural (*six3/6*, *dlx*, *otx*, *pax6* and *nk2.1*), foregut (*foxa* and *gsc*), germline (*nanos*), hindgut/posterior (*bra*, *cdx*, *evx* and *wnt1*), endodermal (*gata456*) and mesodermal (*twist*, *foxc* and *foxf*) markers.

**Figure 9.**
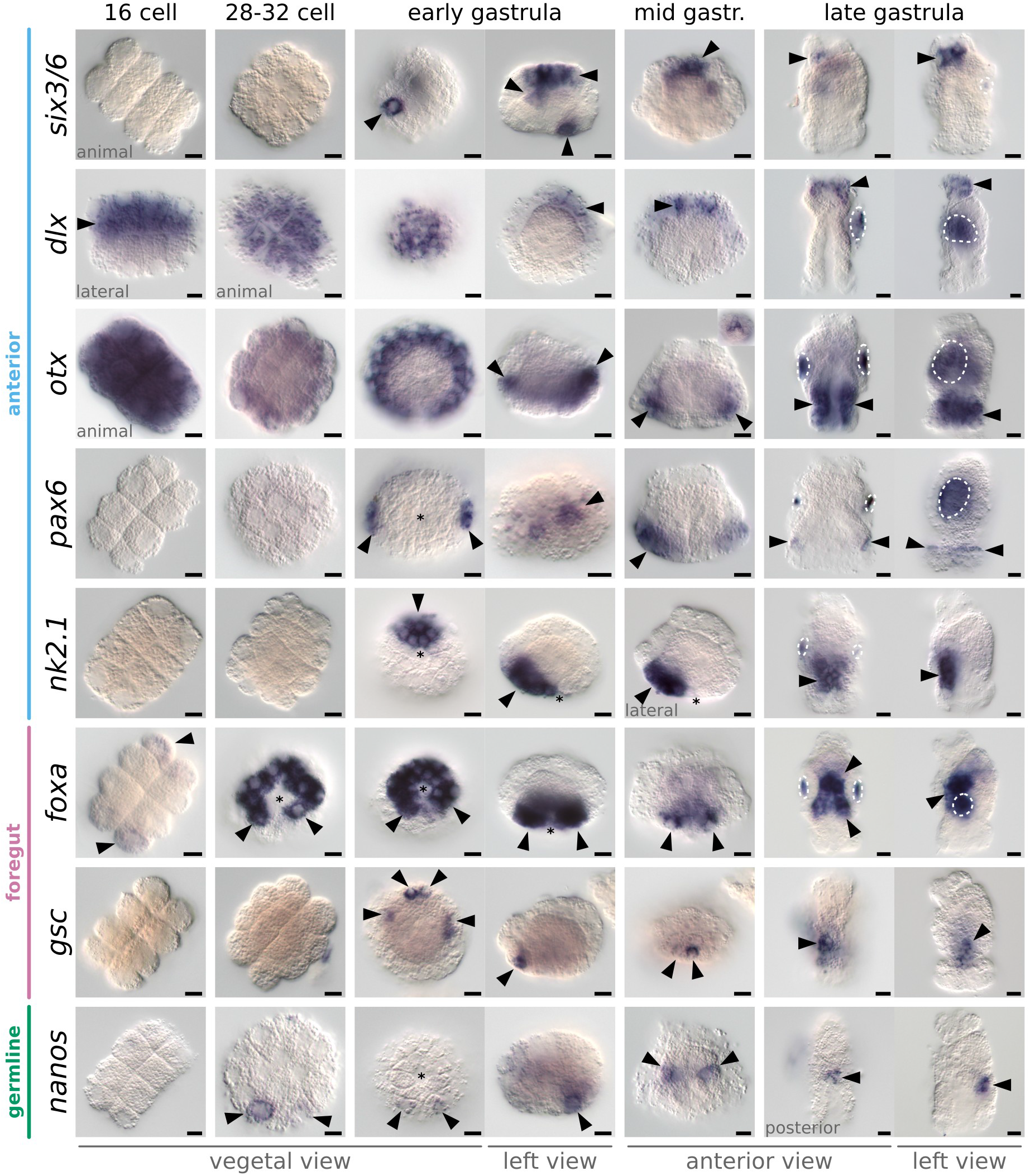
In situ hybridization of anterior, foregut and germline markers during *M. membranacea* embryonic development. Orientation of the embryos is listed below each column and exceptions are marked on individual panels. On vegetal views, B quadrant is top and D quadrant is bottom. On left views anterior (B quadrant) is to the left. All views, except vegetal view, the animal pole is top. Arrowheads indicate expression and dashed areas mark unspecific staining attached to the shell valves of some embryos. Asterisks indicate the position of the blastopore. Scale bar = 10 µm.

**Figure 10.**
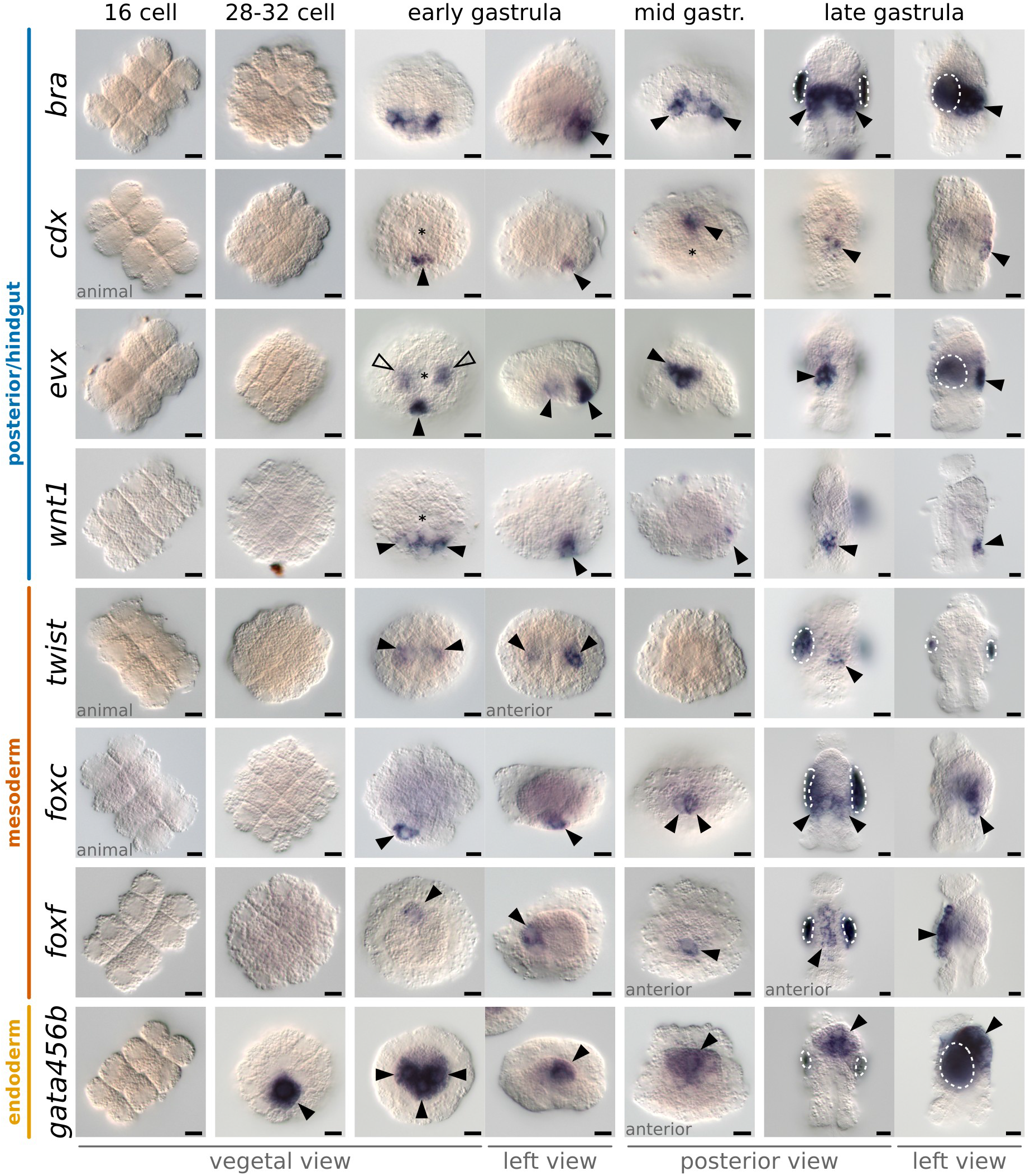
In situ hybridization of posterior/hindgut, mesoderm and endoderm markers in the development of *M. membranacea*. Orientation of the embryos is listed below each column and exceptions are marked on individual panels. On vegetal views, B quadrant is top and D quadrant is bottom. On left views anterior (B quadrant) is to the left. All views, except vegetal view, the animal pole is top. Arrowheads indicate expression and dashed areas mark unspecific staining attached to the shell valves of some embryos. Asterisks indicate the position of the blastopore. Scale bar = 10 µm.

**Figure 11.**
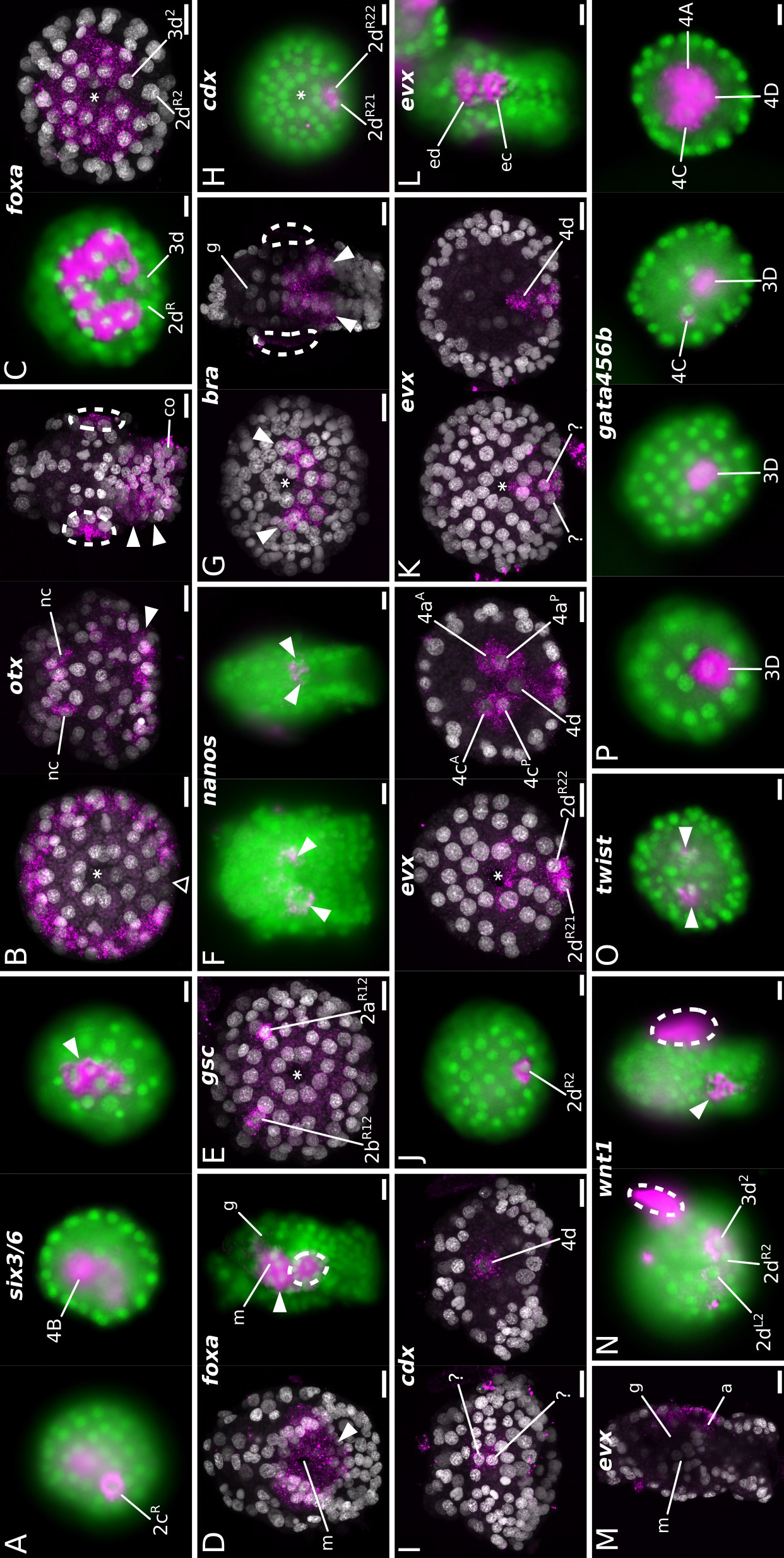
Gene expression details with cell resolution in *M. membranacea*. Selected embryosfrom the in situ hybridizations shown in Figure 9 and 10 observed under a fluorescent lamp (green=nuclei, magenta=signal) or maximum intensity projections from confocal microscopy (gray=nuclei magenta=signal). Arrowheads are pointing to the relevant areas of gene expression while dashed areas mark unspecific background staining. Asterisks mark the position of the blastopore. (A) Expression of *six3/6* at different focal levels. (B) Expression of *otx* in a vegetal view (left) showing the posterior gap in expression (triangle outline), neural cells at the apical disc (nc) of a mid-gastrula embryo and the wider expression in the late gastrula (arrowheads). Expression of *foxa* at the 90-cell stage (left) without signal on the posterior cells 2d^R^ and 3d, and the same posterior gap one cell division cycle later (right). (D) Late gastrula stage (left) viewed from the posterior vegetal end to show the mouth opening with surrounding expression of *foxa* (B quadrant is bottom). On the right, a left side view with *foxa* expression in the mouth region. (E) Bilateral anterior cells expressing *gsc*. (F) Two *nanos*-positive cells during mid-gastrula (left) and late gastrula (right). (G) Posterior and lateral cells on the vegetal ectoderm expressing *bra* (left) and a posterior view of a late gastrula depicting the domain in the posterior epithelium of the vestibule (right). (H) Vegetal view of early gastrula with the two vegetal cells with *cdx* expression (2d^R21^ and 2d^R22^). (I) *cdx* expression observed in two cells at the posterior ectoderm (left), and in the 4d cell (right) at mid-gastrula with two. (J) Expression of *evx* in one posterior ectodermal cell (2d^R2^) on the vegetal side during early gastrulation (left). Progeny of 2d^R2^ expresses *evx* (center) as well as the derivatives of 4a and 4c and the 4d cell. (K) Mid-gastrula stage with *evx* expression in at least two posterior ectodermal cells (left) and in the 4d (right). (L) Posterior view of a late gastrula with *evx* expressed in the posterior endoderm (ed) and ectoderm (ec). (M) Left side view of *evx* expression at the late gastrula with posterior endodermal and ectodermal domains. (N) Expression of *wnt1* during early gastrulation is restricted to three cells, 2d^L2^, 2d^R2^ and 3d^2^ (left) and a posterior cluster of cells at the late gastrula (right). (O) *twist* expression in internalized blastomeres. (P) Expression of *gata456b* from 32-cell stage until early gastrulation. Transcripts are restricted to the 3D until the internalization of vegetal blastomeres, when 4A and 4C initiate the expression of *gata456b*. Scale bar = 10 µm.

We first detected transcripts of *six3/6* in *M. membranacea*, a transcription factor associated to anterior ectodermal patterning (Lowe et al., 2003; Sinigaglia et al., 2013; Steinmetz et al., 2010), during early gastrulation in one outer lateral vegetal plate cell (2c^R2^), one anterior endomesodermal cell (4B) and in 5 cells of the apical disc (Figure 9; Figure 11A). Expression of *six3/6* clears from 2c^R2^ and 4B, but persists in the inner cells of the forming apical organ, a central neural region occupied by serotonergic-positive cells in other cyphonautes larvae (Nielsen and Worsaae, 2010). We detected *dlx* transcripts, a gene involved in neurogenesis and proximodistal patterning (Panganiban and Rubenstein, 2002), in the 8 animal pole cells of the 16 cell stage (1q), broadly in the apical disc during gastrulation and elongation and, finally, localized to the whole apical organ in the late gastrula (Figure 9).

The gene *otx* is involved in anterior ectodermal patterning (Arendt et al., 2001; Boncinelli et al., 1993; Lowe et al., 2003; Marlow et al., 2014; Martín-Durán et al., 2012; Steinmetz et al., 2010; Steinmetz et al., 2011; Umesono et al., 1999) and endomesoderm specification (Harada et al., 2000; Hinman et al., 2003; Mitsunaga-Nakatsubo et al., 2003). In *M. membranacea otx* is expressed in all blastomeres between 2–8-cell stage and gets restricted to the apical octet of the 16-cell stage (Figure 9). At the 32-cell stage, *otx* transcripts localize to the 1q^2^ octet and during gastrulation there are 3 rows of cells expressing *otx* with a posterior gap (Figure 11B). During mid-gastrula two cells in the apical organ express transcripts of *otx* (Figure 11B). In the late larva, *otx* is expressed in the corona and vestibule epithelium (Figure 11B). Expression of *pax6* is first detected during gastrulation, in bilateral patches of the apical ectoderm, and remains as a thin line of expression encircling the embryo above the corona (Figure 9). The gene *nk2.1* is involved in the patterning of the neural plate in vertebrates (Shimamura et al., 1995) and is expressed in anterior and ventral territories including the apical/neural plate and anterior endoderm (Lowe et al., 2003; Marlow et al., 2014; Takacs et al., 2004; Venkatesh et al., 1999). Transcripts of *nk2.1* are present in the progeny of the vegetal cells 2b and 3b in the early gastrula stage (Figure 9). These cells occupy an anterior vegetal position abutting the anterior blastopore lip until the edge of the vegetal plate. After the invagination of the vegetal plate, *nk2.1*-positive cells are lining the anterior portion of the preoral funnel, next to the mouth.

Expression of *foxa* is related to endoderm specification and commonly associated with the blastopore lip and foregut (Arenas-Mena, 2006; Boyle and Seaver, 2010; Oliveri et al., 2006). At the 16-cell stage, we detected faint expression of *foxa* in the outer vegetal blastomeres and in 10 (out of 12) cells surrounding the 4 large blastomeres at the 32-cell stage (2q and 3q, except posterior cells 2d^L^ and 2d^R^) (Figure 9; Figure 11C). Expression of *foxa* persisted in the daughter cells of the next division forming 2 rows of cells around the blastopore with a gap at the posterior end (Figure 11C). With the invagination of the vegetal plate, this region occupies an anterior/lateral position in the vestibule wall, surrounding the mouth region of the late gastrula (Figure 9; Figure 11D). We only found transcripts of *gsc* at the early gastrula stage in two anterior and a bilateral pair of cells at the vegetal plate (Figure 9; Figure 11E). In the late gastrula, *gsc* is expressed in bilateral domains of the vestibule wall which fuse anteriorly.

The germline marker *nanos* (Extavour and Akam, 2003; Juliano et al., 2010) is expressed in two posterior cells of the vegetal plate at the 32-cell stage (2d^L^ and 3d) (Figure 9). In subsequent stages, *nanos* continues restricted to two cells at the posterior portion of the vegetal plate, localizing to the internal sac region of the cyphonautes larva (Figure 11F).

All posterior/hindgut and mesodermal markers only initiate expression during gastrulation. The gene *bra* can have multiple roles, but it is generally related to mesoderm and posterior/hindgut patterning (Technau, 2001). Expression of *M. membranacea bra* in the early gastrula occurs at the vegetal plate in a posterior band of cells near the blastopore lip (Figure 10; Figure 11G). It localizes to 6–8 cells at the posterior end of the mid gastrula and a broad portion of the posterior and lateral vestibule ectoderm (Figure 11G). *M. membranacea bra* expression domain reaches the posterior portion of the preoral funnel as well as the future hindgut area of the larva (Figure 10). A single posterior vegetal plate cell (2d^R2^) and its daughter cells (2d^R21^ and 2d^R22^) express the posterior/hindgut markers *cdx* and *evx* at the early gastrula (Figure 10; Figure 11H; Figure 11J). During gastrulation, *cdx* and *evx* continue to be expressed at the posterior edge of the vegetal plate (Figure 10; Figure 11J; Figure 11L) and localize to the posterior vestibule ectoderm (hindgut) of the late gastrula (Figure 10; Figure 11L). At this stage, *evx* is also found in the posterior region of the gut (Figure 11L; Figure 11M). We also detected a transient *evx* expression in the two internalized blastomeres 4a and 4c of the early gastrula. Finally, *wnt1* is expressed in a row of 3–5 cells (including 2d^L2^, 2d^R2^ and 3d^2^) posterior to the blastopore during gastrulation (Figure 10; Figure 11N). At the late gastrula, *wnt1* is detected at the posterior-most vestibule ectoderm, positioned between the corona and hindgut (Figure 10; Figure 11N).

Expression of *twist*, a central regulator in mesoderm differentiation (Technau and Scholz, 2003), only occurs in a portion of embryos during the development of *M. membranacea*. We detected colorimetric signal in bilateral internalized cells of the early gastrula – possibly 4a, 4c or derivatives – as well as at the anterior end of the late gastrula (Figure 10; Figure 11O). Transcripts of *foxc*, commonly expressed in anterior and posterior mesodermal domains (Häcker et al., 1995; Passamaneck et al., 2015; Shimeld et al., 2010), are present in one unidentified posterior vegetal plate cell of the early gastrula and two similarly positioned cells during mid gastrulation (Figure 10). In the late gastrula, *foxc* expression is located in the internal sac area. The gene *foxf* is a transcription factor involved in mesoderm patterning and expressed mainly in visceral and anterior territories (Mazet et al., 2006; Passamaneck et al., 2015; Pérez Sánchez et al., 2002; Shimeld et al., 2010; Zaffran et al., 2001). In *M. membranacea* it is expressed in the mesodermal cell 4b in the early and mid gastrula stages (Figure 10). This cell and its descendants divide subsequently from basal to apical, forming a distinct frontal row of mesodermal cells expressing *foxf* at the anterior portion of the late gastrula.

We found two copies of the endomesodermal marker *gata456* (Patient and McGhee, 2002) in the transcriptome of *M. membranacea*. While the gene *gata456a* is not expressed at detectable levels in any of the analyzed stages, *gata456b* is strongly expressed in endodermal tissues of the bryozoan. The expression of *gata456b* initiates early, in the vegetal 3D blastomere at the 32-cell stage (Figure 10; Figure 11P). The expression expands to adjacent lateral blastomeres 4A and 4C in the early gastrula, and in subsequent stages *gata456b* continues to be expressed in the endodermal tissues forming the gut of the cyphonautes larva (Figure 10; Figure 11P).

## Discussion

The phylogenetic position of bryozoans provides a valuable opportunity to investigate the evolution of developmental traits. Even though the kinship of Bryozoa remains inconclusive – the group is either related to Entoprocta and Cycliophora (e.g. Hejnol et al., 2009) or to Phoronida and Brachiopoda (e.g. Nesnidal et al., 2013), or both (Kocot et al., 2016) – most phylogenetic analyses place the bryozoans nested within the Spiralia (Kocot, 2016; Kocot et al., 2016; Laumer et al., 2015; Nesnidal et al., 2013; Struck et al., 2014). This indicates that the ancestral cleavage pattern of Spiralia – spiral cleavage – must have been modified in the bryozoan lineage during evolution (Hejnol, 2010). Hereby we examine the similarities and differences between the embryogeneses of the bryozoan *M. membranacea* and that of spiral-cleaving embryos, by integrating cell lineage and molecular data, and provide a hypothesis for the evolution of bryozoan development from a spiral-cleaving ancestor.

### Specification of the D quadrant

One critical event of animal embryogenesis is the establishment of the dorsoventral polarity. In spiral-cleaving embryos this event is tied to the specification of the D quadrant during development (Freeman and Lundelius, 1992). In species where the first two embryonic cell divisions are unequal, the D quadrant is determined early by the asymmetric distribution of maternal cytoplasmic determinants, while in species that form equal-sized blastomeres at the 4-cell stage, the D quadrant is specified at the 32-cell stage by inductive interactions mediated by cell contacts between micromeres and macromeres (Biggelaar, 1977; Freeman and Lundelius, 1992; Gonzales et al., 2006; Martindale, 1986; Martindale et al., 1985). In the current work, we found evidence that the specification of the D quadrant in the equal, biradial-cleaving bryozoan *M. membranacea* resembles that of equal, spiral-cleaving molluscs in the timing of specification, pattern of MAPK activation and asynchrony of the D quadrant cell divisions post-specification.

In equal-cleaving molluscs, the specification of the D quadrant correlates with the activation of the MAPK pathway in the 3D macromere only (Koop et al., 2007; Lambert and Nagy, 2003). In *M. membranacea*, whose equal-sized blastomeres at the 4-cell stage give rise to perfectly symmetrical embryonic quadrants, that are indistinguishable from each other until gastrulation, the earliest molecular asymmetry we could detect is the activation of the MAPK pathway in a single vegetal blastomere that originates the posterior portion of the larval body. As in equal-cleaving molluscs, MAPK is activated in the bryozoan 3D blastomere on the 5th round of cell divisions, suggesting the D quadrant of *M. membranacea* is specified as early as the 28-cell stage. This might indicate that bryozoans and equal-cleaving molluscs undergo similar developmental mechanisms of D quadrant specification (but see below). Interestingly, most equal-cleaving spiralians studied so far exhibit a single MAPK-activated blastomere during early development, while unequal-cleaving species show diverse deviating patterns (see Table 1), thus suggesting that this pattern of MAPK activity is a common feature of equal-cleaving embryogenesis independent of its cleavage pattern.

**Table 1.** MAPK activity in spiralians.

| taxon | species | ABCD sizes | MAPK activity | MAPK inhibition (U0126) |
| --- | --- | --- | --- | --- |
| Mollusca (gastropod) | *Ilyanassa obsoleta* | Unequal | Strong signal in the cytoplasm around nucleus of the 3D cell. Activity spread on the overlying 2d1 and 2d2; then 1c12, 1a12, 1b12; 4d cell, but not 4D; derivatives of the second and third quartet forming an arc centered in the dorsal midline; micromeres 2b12 and 3b1 transiently; undetectable 6h after 4d formation; never detected in 3A, 3B, 3C, 2b2 or 3a (Lambert and Nagy, 2001). | 10μM at 23°C – Disorganized and incomplete larvae. MAPK activation is required on the micromeres themselves, not only in 3D. Alters the cleavage pattern in the third quartet of micromeres and timing of division of the 4d (Lambert and Nagy, 2001). |
| Mollusca (gastropod) | *Crepidula fornicata* | Equal-like | 1a1-1d1 micromeres, fainter in 1a2-1d2; 3D cell, persisting in 4d; also 3d1-3d2 and 1a12-1d12 (Henry and Perry, 2007). | 10–25μM at 20–24°C – Radialized embryos with no differentiated axial properties or cell fates (Henry and Perry, 2007). |
| Mollusca (gastropod) | *Stagnicola palustris* (= *Lymnaea palustris*)* | Equal | 3D only (Lambert and Nagy, 2003). | - |
| Mollusca (gastropod) | *Lottia scutum* (= *Tectura scutum*)* | Equal | 3D only (Lambert and Nagy, 2003). | 25–50μM at 14°C – Prevented normal development of micromeres producing the foot, the shell and affecting the differentiation of eyes (Lambert and Nagy, 2003). |
| Mollusca (gastropod) | *Haliotis asinina* | Equal | 3D only (Koop et al., 2007). | 10–50μM at 25°C – Retarded development and failure to develop larval musculature, shell and foot. But gene expression not completely radialized and treated embryos at lower concentrations were only mildly abnormal (Koop et al., 2007). |
| Mollusca (gastropod) | *Patella vulgata* | Equal | 3D only (Lartillot et al., 2002a). | 10–50μM at ~14°C – Overall morphology somewhat disturbed, but without gross axial or gastrulation defects (Lartillot et al., 2002a). |
| Mollusca (polyplacophoran) | *Chaetopleura apiculata* | Equal | 3D only (Lambert and Nagy, 2003). | - |
| Annelida (sedentary polychaete) | *Hydroides dianthus* (= *Hydroides hexagonus*)* | Equal | 4d only (Lambert and Nagy, 2003). | - |
| Annelida (sedentary polychaete) | *Capitella teleta* | Unequal | Cells around the blastopore during gastrulation (Amiel et al., 2013). | 5–50μM at 14°C – Compact shaped larva with less muscles and disorganized axon tracts, but without affecting the D quadrant specification, organizing activity or establishment of larval body axes (Amiel et al., 2013). |

| **taxon** | **species** | **ABCD sizes** | **MAPK activity** | **MAPK inhibition (U0126)** |
| --- | --- | --- | --- | --- |
| Annelida (errant polychaete) | *Platynereis dumerilii* | Unequal | Two dorsal cells (nephroblasts) and micromeres around the blastopore (Pfeifer et al., 2014). | 10–50µM at 18°C – Shortened overall morphology with disorganized muscle and neural tracts (Pfeifer et al., 2014). |
| Annelida (errant polychaete) | *Alitta virens* | Unequal | 8-cell stage at 1c, 1d and 1D; all micromeres at 16-cell (no vegetal activity); 3d and 3D activity, off in micromeres; then 1q1 and 2d descendant, 3c, 3d, 3D. Fades after that (Kozin et al., 2016). | 40µM at 14°C – Motile larvae with normal bilateral symmetry, mesoderm present but different phenotypes (Kozin et al., 2016). |
| Bryozoa (gymnolaemate) | *Membranipora membranacea* | Equal | 3D only. | 1–25µM at 10–15°C – Delayed shortened larvae. |

\*Synonyms taken from WoRMS database.

The developmental role of the MAPK pathway in spiralians, however, is far less clear. Drug treatments to block MAPK activity in molluscs cause radialized larvae that lack muscles, shell and foot, suggesting that the MAPK pathway is involved in the signaling underlying the D quadrant specification (Henry and Perry, 2007; Koop et al., 2007; Lambert and Nagy, 2001; Lambert and Nagy, 2003). While we found severely disturbed *M. membranacea* embryos in our MEK inhibitor experiments, the abnormal cleavage leading to the severe phenotype occurs before any detectable MAPK activity, as revealed by the 4D recordings. Thus, the developmental defect we observe in the severe phenotype is unlikely to be a direct consequence of the inhibition of MAPK signaling in the 3D blastomere, unless MAPK signaling is active in undetectable levels at the 16-cell stage. Instead, most of our experiments indicate that the inhibition of the MAPK pathway does not cause dorsoventral defects in *M. membranacea*. In fact, when treated with the same MEK inhibitor concentration we used in the bryozoan, equal-cleaving molluscs still show a latent dorsoventral polarity or develop without axial defects (Koop et al., 2007; Lartillot et al., 2002a), which might indicate that MAPK is not the sole signaling agent for the D quadrant specification in spiralians.

Once the D quadrant has been determined, it typically shows asynchronous cell divisions in relation to the other quadrants of spiral-cleaving embryos (Guralnick and Lindberg, 2001). For instance, the 3D macromere in the mollusc *Patella vulgata* (Biggelaar, 1977) and the 1d derivatives of *Ilyanassa obsoleta* (Clement, 1952; Goulding, 2009) undergo a late division. Our analyses of *M. membranacea* cell lineage indicate similar asynchronous cell divisions in the D quadrant, which include the 3D blastomere and 1d derivatives of the bryozoan. Therefore, the specification of the D quadrant seems to be correlated with subsequent changes in the cell cycle timing in both *M. membranacea* and spiral-cleaving embryos.

Overall, *M. membranacea* exhibits a similar pattern and timing of MAPK activation, as well as equivalent asynchronous cell divisions in the D quadrant, when compared to equal-cleaving molluscs. Given the phylogenetic position of bryozoans, these similarities might suggest that some of the underlying traits of spiral-cleaving embryos were maintained during the evolutionary transition from spiral to biradial cleavage. The comparison also reveals that equal cleavage might be associated with a single D quadrant MAPK-activated blastomere in spiralian development. Nevertheless, the MAPK pathway is still poorly sampled in spiralians, and other spiral and non-spiral-cleaving groups, such as phoronids, nemerteans, polyclads, rotifers and gastrotrichs, need to be investigated to properly understand the roles and the evolution of MAPK signaling in spiralian development.

### Comparative spiralian fate maps

The stereotypic cleavage pattern of spiral cleavage permits to identify putative homologous blastomeres between different spiralians, and to compare their fates in the larval/adult tissues (Conklin, 1897; Mead, 1897; Treadwell, 1901; Wilson, 1892; Wilson, 1898). These studies revealed that homologous early blastomeres share mostly-similar fates in various clades (Hejnol, 2010; Henry and Martindale, 1999; Lambert, 2010; Nielsen, 2004; Nielsen, 2005). The cleavage of *M.membranacea* clearly differs from the spiral cleavage pattern, which complicates the identification of homologous blastomeres between the bryozoan and a spiral-cleaving embryo. However, we established a common developmental feature to base our comparative cell lineage and gene expression analyses. In both spiral and bryozoan embryogenesis, the vegetal blastomeres sequentially give rise to quartets of daughter cells, while remaining at the vegetal-most portion of the embryo until being internalized during gastrulation. We thus compare the quartets of *M. membranacea* to the quartets of spiral-cleaving embryos in terms of gene expression and fate in the larval tissues. We find the quartets have a similar molecular identity and contribute to the same set of structures in the larvae of bryozoan and spiral-cleaving groups, and that the subset of blastomeres that gives rise to these structures partially overlap (Figure 12). This indicates that bryozoans might share a common embryonic patterning of early blastomere fates with other spiralians, and that, in the current phylogenetic scenario, such developmental trait has remained conserved despite the drastic modification in the cleavage pattern from spiral to biradial.

**Figure 12.**
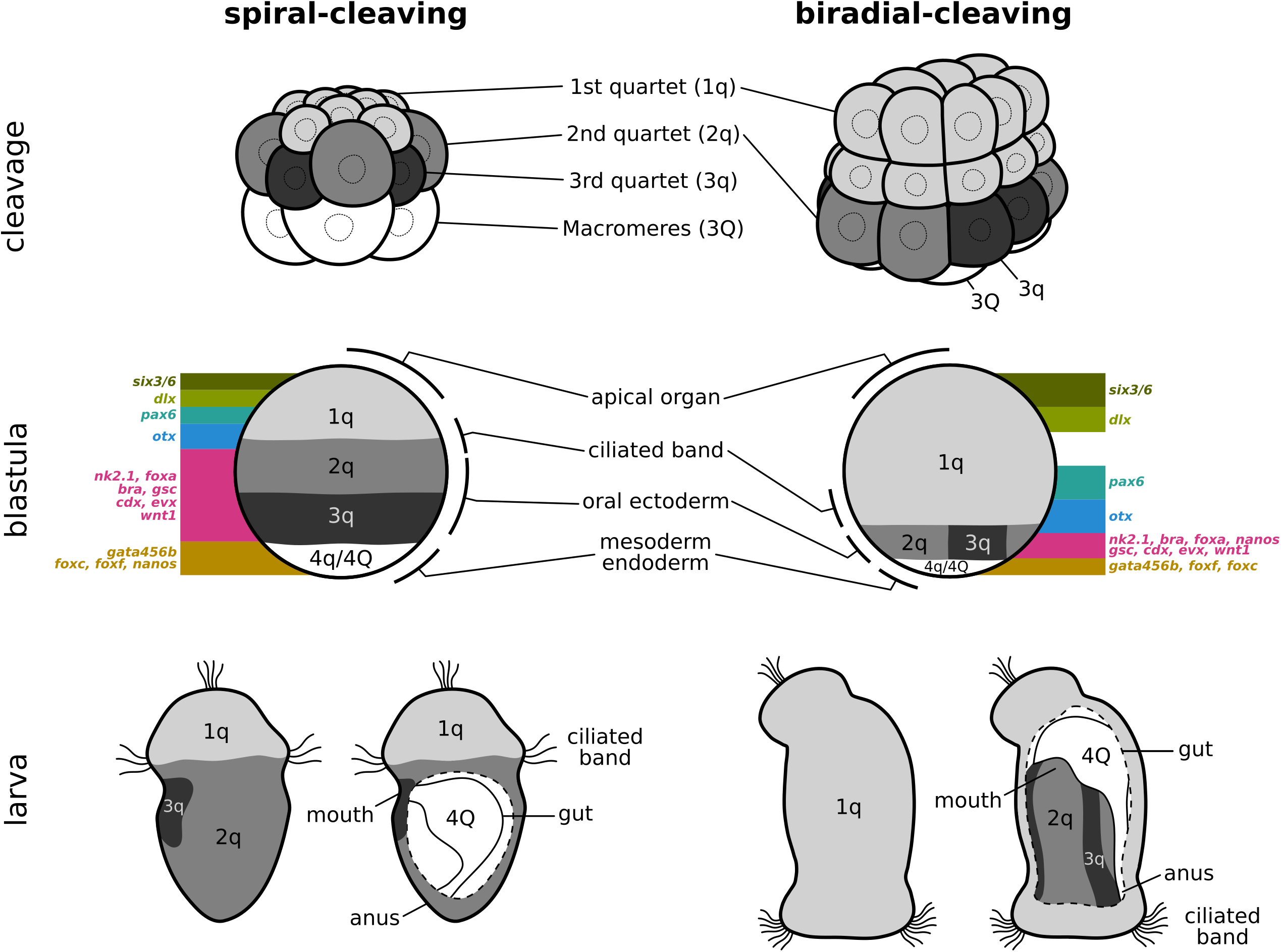
Bryozoan development in comparison to spiral cleavage. Line drawings represent cleavage, blastula and larval stages of a generalized spiral-cleaving embryo and the biradial-cleaving embryo of the bryozoan *M. membranacea*. Shades of grey indicate the first, second and third quartets and their respective fates in the blastula and larval tissues. The fourth quartet, macromeres and descendants are depicted in white. A simplified summary of the gene expression domains is mapped to the blastula stage.

#### First quartet: apical organ and ciliated band

The first quartet of micromeres in spiral-cleaving embryos contributes to the apical organ, the ciliated band and all the ectoderm in between (Ackermann et al., 2005; Boyer et al., 1998; Child, 1900; Conklin, 1897; Damen and Dictus, 1994; Dictus and Damen, 1997; Hejnol et al., 2007; Henry and Martindale, 1998; Henry et al., 2004; Maslakova et al., 2004; Meyer et al., 2010; Wilson, 1892). In the *M. membranacea*, the first quartet of animal blastomeres also gives rise to these ectodermal structures of the cyphonautes larva. This suggests that in both the bryozoan and spiral-cleaving embryos the third cleavage demarcates a split in the embryonic fate map, in which the first quartet of animal blastomeres only gives rise to the ectodermal structures placed towards the animal pole, while the progeny of the vegetal blastomeres (i.e. the 2nd, 3rd and 4th quartets) originates a different set of larval structures (see the next sections). When we compare in more detail the specific fates of the descendants of the first quartet, we find that some blastomeres contributing to the apical organ or ciliated band of the cyphonautes larva indeed contribute to the respective structures of spiral-cleaving larvae – but that this similarity is not complete, and different subsets of blastomeres contribute to the apical organ and ciliated band.

The apical organ, for example, is usually formed by the progeny of the apical-most 1q^1^ micromeres in groups with spiral cleavage (Nielsen, 2004; Nielsen, 2005). While the apical organ of *M. membranacea* larva is also derived from the apical-most subset of 1q_i_^1^ (=1q^11^), descendants of 1q_e_^1^ (=1q^21^) contribute to the structure as well. We find a similar situation when comparing the embryonic origin of the corona (i.e. the ciliated band of the cyphonautes larva) with the prototroch – a ciliated band considered to be an ancestral trait for the larval stages of trochozoan spiralians (Rouse, 1999). The prototroch of annelids and molluscs is formed by 1q (accessory and primary trochoblasts) and 2a–c (secondary trochoblasts) descendants (Damen and Dictus, 1994; Hejnol et al., 2007; Henry et al., 2007). Our data reveals that the corona of *M.membranacea* is formed by blastomeres equivalent to the accessory/primary trochoblasts of the prototroch (Damen and Dictus, 1994). But, unlike spiral-cleaving embryos, the second quartet does not contribute to the ciliated band of the bryozoan larva (see Figure S7). In general, we find that equivalent early blastomeres of the bryozoan and spiral-cleaving embryos contribute to similar larval structures, but that the fate of the progeny of these early blastomeres only partially overlap between the bryozoan and spiral-cleaving embryos. These observations suggest that during the evolution of the bryozoans, shifts in the blastomere fates occurred in late embryogenesis while the early embryonic patterning, presumably inherited from a spiral cleavage ancestor, might have remained conserved.

The bryozoan fate map indicates that the apical organ, outer ectoderm and corona of the cyphonautes larva have a similar embryonic origin than the apical organ, pretrochal elements and prototroch of spiral-cleaving embryos, respectively. They all derive from the first quartet of blastomeres. The fate map similarity is paralleled by molecular data, since the genes expressed in this region of *M. membranacea* have an equivalent spatial arrangement in other spiralian embryos. The expression of *six3/6*, *dlx* and *otx* in the ectoderm of *M. membranacea* also reflects the conserved staggered expression of regulatory transcription factors along the animal-vegetal axis of metazoan embryos (Gonzalez et al., 2016; Lowe et al., 2003; Sinigaglia et al., 2013; Steinmetz et al., 2010). *six3/6* and *dlx* are expressed at the animal end while *otx* is expressed in the basal-most progeny of the first quartet. Other spiralian embryos display a similar arrangement of these transcripts (Arendt et al., 2001; Hiebert and Maslakova, 2015a; Hiebert and Maslakova, 2015b; Marlow et al., 2014; Martín-Durán et al., 2015; Perry et al., 2015; Pineda and Saló, 2002; Santagata et al., 2012; Steinmetz et al., 2010; Steinmetz et al., 2011) (also see Table 2). Therefore, the region derived from the first quartet of *M. membranacea* match the pretroch region of spiral-cleaving embryos.

**Table 2.** Gene expression patterns in spiralians.

| **taxon** | ***six3/6*** | ***dlx*** | ***otx*** | ***pax6*** | ***nk2.1*** | ***foxa*** | ***gsc*** | ***nanos*** |
| --- | --- | --- | --- | --- | --- | --- | --- | --- |
| Rotifera | - | - | - | anterior bilateral patches (Boell and Bucher, 2008) | - | - | - | oocyte and posterior patch of ovary cells (Smith et al., 2010) |
| Micrognathozoa | - | - | - | - | - | - | - | - |
| Gnathostomulida | - | - | - | - | - | - | - | - |
| Platyhelminthes | brain branches (Pineda and Saló, 2002) | brain, eyes, head and pharynx (Lapan and Reddien, 2011) | brain (Umesono et al., 1999) | eyes (Callaerts et al., 1999) | parenchyma (Garcia-Fernández et al., 1993) | pharynx duct (Martín-Durán et al., 2010) | central nervous system (Martín-Durán and Romero, 2011) | germline and eye precursors (Handberg-Thorsager and Saló, 2007; Sato et al., 2006) |
| Gastrotricha | - | - | - | - | - | - | - | - |
| Mollusca | anterior blastopore lip, apical ganglia and ocelli (Perry et al., 2015) | all animal pole cells and general ectoderm (Lee and Jacobs, 1999) | prototroch (Nederbragt et al., 2002a); blastopore lip, ventral midline, mouth, mesoderm and velum (Perry et al., 2015) | optic region, brain and arms (Tomarev et al., 1997) | anterior most region of the stomodeum, crescent above the mouth (Perry et al., 2015) | blastopore lips, endoderm and mesoderm (Lartillot et al., 2002b; Perry et al., 2015) | anterior mesoderm during gastrulation, foregut and mantle edge in larval stages (Lartillot et al., 2002b); ecto and endomesoderm and anterior of blastopore lip (Perry et al., 2015) | animal pole cells then two posterior bilaterally symmetrical cells (Kranz et al., 2010); or 4d (Rabinowitz et al., 2008) |
| Annelida | apical domain encircling apical organ (Marlow et al., 2014; Steinmetz et al., 2010) | apical ectoderm, apical organ, prototroch and ganglia (McDougall et al., 2011) | early apical organ, foregut and prototroch (Arendt et al., 2001; Marlow et al., 2014; Steinmetz et al., 2010), also brain and posterior growth zone (Boyle et al., 2014) | anterior bilateral patches (Arendt et al., 2002; Denes et al., 2007; Quigley et al., 2007) | apical ventral domain above prototroch (Marlow et al., 2014) and bilateral domains flanking blastopore (Boyle et al., 2014) | vegetal plate ring with cleared posterior expression, foregut and hindgut (Arenas-Mena, 2006; Boyle and Seaver, 2008; Boyle and Seaver, 2010; Boyle et al., 2014) | foregut and bilateral neural domains (Arendt et al., 2001; Boyle et al., 2014) | mesodermal posterior growth zone (Rebscher et al., 2007); anterior ectodermal clusters, foregut, posterior segmental zone (Dill and Seaver, 2008); ectodermal precursor (Kang et al., 2002) |
| Nemertea | apical organ, ciliary band and cephalic imaginal discs (Hiebert and Maslakova, 2015a); anterior invagination and apical disc (Hiebert and Maslakova, 2015b); anterior cephalic imaginal discs, cephalic lobes and mouth (Martín-Durán et al., 2015) | - | around blastopore, anterior imaginal disc, anterior region and gut (Martín-Durán et al., 2015) | - | - | mouth and pharynx, also head and posterior (Martín-Durán et al., 2015) | pair of bilateral clusters near the blastopore and anterolateral in juvenile (Martín-Durán et al., 2015) | - |
| Brachiopoda | apical domain, endoderm (Martín-Durán et al., 2016b; Santagata et al., 2012) | - | apical domain, foregut, endoderm (Martín-Durán et al., 2016b; Passamaneck et al., 2011) | apical lobe (Passamaneck et al., 2011; Vellutini and Hejnos, 2016) | apical domain, foregut (Martín-Durán et al., 2016b; Santagata et al., 2012) | foregut, endoderm (Martín-Durán et al., 2016b) | apical domain, foregut (Martín-Durán et al., 2016b) | - |
| Phoronida | - | - | - | - | - | - | - | - |
| Entoprocta | - | - | - | - | - | - | - | - |
| Bryozoa | apical organ,<br>anterior<br>endoderm and<br>lateral vegetal<br>plate cell | apical organ | corona, apical<br>disc during<br>gastrulation | bilateral domains<br>above the corona | vegetal anterior<br>domain, foregut | vegetal<br>ectoderm<br>around<br>blastopore,<br>foregut | two anterior<br>vegetal cells and<br>pair of bilateral<br>domains | two posterior cells<br>on the vegetal plate |

**Table 2 (continued).** Gene expression patterns in spiralian.
| *taxon* | *bra* | *cdx* | *evx* | *wnt1* | *twist* | *foxc* | *foxf* | *gata456* |
| --- | --- | --- | --- | --- | --- | --- | --- | --- |
| Rotifera | - | - | - | - | - | - | - | - |
| Micrognathozoa | - | - | - | - | - | - | - | - |
| Gnathostomulida | - | - | - | - | - | - | - | - |
| Platyhelminthes | - | - | - | posterior end (Adell et al., 2009; Petersen and Reddien, 2008) | pharynx muscle (Martín-Durán et al., 2010) | - | - | blind gut (Martín-Durán and Romero, 2011) |
| Gastrotricha | - | - | - | - | - | - | - | - |
| Mollusca | 3D and mostly posterior edge of the blastopore (Lartillot et al., 2002a; Perry et al., 2015); posterior of blastopore (Koop et al., 2007) | posterior ectoderm (Fritsch et al., 2016; Le Gouar et al., 2003; Perry et al., 2015), mesoderm close to the blastopore (Le Gouar et al., 2003; Perry et al., 2015) and mouth/esophagus (Perry et al., 2015) | - | - | anterior mesoderm but not in its mesenchymal progeny (Nederbragt et al., 2002b; Perry et al., 2015) | mantle, foregut and trunk mesoderm (Shimeld et al., 2010) | bilateral mesodermal cells next to endoderm (Shimeld et al., 2010) | - |
| Annelida | A-D macromeres, foregut and hindgut (Arendt et al., 2001; Boyle et al., 2014) | hindgut, posterior end (Fröbuis and Seaver, 2006; Hui et al., 2009; Kulakova et al., 2008; Rosa et al., 2005; Samadi and Steiner, 2010); 4d-derived bilaterally symmetric lateral cells and early segmental mesoderm (Fröbuis and Seaver, 2006) | posterior and bilateral bands that can be internalized (Rosa et al., 2005; Seaver et al., 2012); teloblasts and ventral nerve cord (Seaver et al., 2012; Song et al., 2002) | posterior end (Prud'homme et al., 2003; Seaver and Kaneshige, 2006) | mesoderm derivatives during larval development (Dill et al., 2007; Pfeifer et al., 2013) | head and and foregut mesoderm (Shimeld et al., 2010) | brain, foregut and posterior trunk mesoderm (Shimeld et al., 2010) | endo/mesodermal precursors 2Q, 4Q and 4a-c but not 4d (Boyle and Seaver, 2008; Boyle and Seaver, 2010; Wong and Arenas-Mena, 2016), endomesoderm and paired lateral bands (Gillis et al., 2007) |
| Nemertea | - | posterior end then subepidermal (Hiebert and Maslakova, 2015b), posterior end and posterior endoderm (Martín-Durán et al., 2015) | posterior end and scattered cells in juvenile (Martín-Durán et al., 2015) | - | larval imaginal discs and broadly later (Martín-Durán et al., 2015) | - | - | larval blind gut and juvenile gut (Martín-Durán et al., 2015) |
| Brachiopoda | blastopore, foregut (Martín-Durán et al., 2016b) | posterior ectoderm and hindgut (Altenburger et al., 2011; Martín-Durán et al., 2016b) | hindgut, posterior ectoderm (Martín-Durán et al., 2016b) | posterior end of blastopore (Vellutini and Hejnal, 2016) | mesoderm (Martín-Durán et al., 2016b; Passamaneck et al., 2015) | apical and pedicle mesoderm, neuroectoderm (Martín-Durán et al., 2016b; Passamaneck et al., 2015) | anterior mesoderm (Martín-Durán et al., 2016b; Passamaneck et al., 2015) | endoderm and posterior mesoderm (Martín-Durán et al., 2016b; Passamaneck et al., 2015) |
| Phoronida | - | - | - | - | - | - | - | - |
| Entoprocta | - | - | - | - | - | - | - | - |
| Bryozoa | posterior<br>ectoderm,<br>larval hindgut | posterior<br>ectoderm,<br>larval<br>hindgut | 2dr2 posterior<br>ectodermal<br>cell and<br>progeny, 4a,<br>4c, and 4d<br>mesodermal<br>blastomeres,<br>larval hindgut | posterior end | bilateral<br>mesodermal<br>blastomeres | posterior<br>ectodermal cell,<br>internal sac | anterior larval<br>mesoderm | posterior vegetal<br>blastomeres, larval<br>gut |

In *M. membranacea* the blastomeres that form the apical organ express *six3/6* and *dlx* from the 16-cell stage, suggesting that these genes might be involved in the establishment of the embryonic animal/vegetal identities, and possibly in the molecular patterning of the cyphonautes apical organ. The expression of *otx* in the bryozoan is associated to the corona, the ciliated band of the cyphonautes larva, similar to other spiralians where *otx* is expressed near or in the larval ciliated band (Arendt et al., 2001; Nederbragt et al., 2002a; Steinmetz et al., 2011). The gene is an interesting example because it provides the opportunity to integrate the cell lineage and gene expression data between the bryozoan and spiral-cleaving embryos. As explained above, the ciliated band of trochophore larvae, the prototroch, is formed by the contribution of first quartet and second quartet blastomeres (Damen and Dictus, 1994; Henry et al., 2007), while the corona of *M.membranacea* derives solely from first quartet blastomeres, which are the putatively homologous to the primary trochoblasts of the prototroch. In the mollusc *Patella vulgata*, *otx* is expressed in all prototroch cells (Nederbragt et al., 2002a). Interestingly, the second quartet blastomeres of *M.membranacea* – the set of blastomeres that form the secondary trochoblasts in the prototroch – also express *otx*, as observed in *P. vulgata*, even though these cells do not contribute to the corona of the cyphonautes larva. This observation suggests that presumptive homologous blastomeres between the bryozoan and other spiralians might still share a similar molecular identity, even though they do not form similar tissues.

Overall, our work reveals that the first quartet of *M. membranacea* embryo and the first quartet of spiral-cleaving embryos originate a similar set of larval structures, and give rise to a larval body region with similar molecular profile. Thus, the outer ectodermal region of the cyphonautes larva corresponds, in developmental terms, to the head region of other spiralians.

#### Second and third quartet: mouth

The second and third blastomere quartets of spiral-cleaving embryos contribute to a diverse set of ectodermal structures, such as the foregut, ciliated bands, neurons, the mollusc shell gland and foot, the annelid trunk and nerve cord, as well as ecto-mesodermal muscle cells (Ackermann et al., 2005; Boyer et al., 1998; Dictus and Damen, 1997; Hejnol et al., 2007; Henry and Martindale, 1998; Henry et al., 2004; Lyons et al., 2015; Maslakova et al., 2004; Meyer et al., 2010; Render, 1997). In *M. membranacea* these blastomeres form the whole vegetal ectoderm that gives rise to the vestibule epithelium, including the preoral funnel and posterior ectoderm of the cyphonautes larva. In most spiralians the second and third quartets are the blastomeres surrounding the blastopore – the orifice formed at the site of endomesoderm internalization (Lankester, 1877), whose developmental fate has been a significant trait for the discussions about metazoan evolution (Hejnol and Martín-Durán, 2015; Martindale and Hejnol, 2009; Martín-Durán et al., 2016b). Nevertheless, the fate of the blastopore in bryozoans still remains open to discussion (Gruhl, 2009; Marcus, 1938; Prouho, 1892; Zimmer, 1997).

Even though in most gymnolaemate bryozoans the blastopore closes after gastrulation (Pace, 1906; Prouho, 1892), or in some cases, an orifice is not formed at all (Calvet, 1900), an ultra-structural study in *M. membranacea* revealed that its blastopore remains open until the larval stage (Gruhl, 2009). Our cell lineage data indicate that cells at the blastopore lip give rise to the preoral funnel of *M. membranacea*, and that the endodermal cells lining the blastopore form the anterior portion of the larval gut. We also found the foregut marker *foxa* is expressed around most of the blastopore lip, except for a couple of posterior rows, and that *foxa* expression persists around the future larval mouth, indicating that most cells associated with the blastopore of *M.membranacea* have a foregut molecular identity. Thus, independent ultrastructural, molecular and cell lineage data provides robust evidence for a persistent blastopore and the protostomic development of *M. membranacea*, as previously suggested (Gruhl, 2009).

We found that the vegetal ectoderm – the cells derived from the second and third quartet – exhibits an anteroposterior polarity, as revealed by the differential expression of molecular markers. The anterior/foregut markers *nk2.1*, *foxa* and *gsc* are expressed in a region opposed to posterior/hindgut markers *bra*, *cdx*, *evx* and *wnt1*, which are generally restricted to the D quadrant. Transcripts of *nk2.1* are restricted to the B quadrant in a comparable position, in relation to the cyphonautes anteroposterior axis, to the anterior/ventral expression found in other bilaterians (Lowe et al., 2003; Marlow et al., 2014; Martín-Durán et al., 2016b; Perry et al., 2015; Tessmar-Raible et al., 2007). In a similar fashion, transcripts of *wnt1*, a gene commonly expressed at the posterior end of bilaterians (Bolognesi et al., 2008; Holland et al., 2000; Pani et al., 2012; Prud’homme et al., 2003), occur at the posterior region of the vegetal ectoderm of *M. mem-branacea* (see Table 2 for a broader comparison of gene expression within spiralians). This suggests that at least some molecular aspects of the axial patterning remain conserved in the cyphonautes larva.

In some cases, the transcripts of *M. membranacea* are not only located at a similar position along the anteroposterior axis, but also in the putative homologous blastomeres to spiral-cleaving embryos. An example is the expression of *foxa* between the bryozoan and the annelid *Hydroideselegans* (Arenas-Mena, 2006). In both, *foxa* is expressed in the second quartet blastomeres early in development, and in the cells that surround the blastopore during gastrulation, with a peculiar posterior gap (Arenas-Mena, 2006). Another comparable cellular expression is the gene *bra*, expressed in the second and third quartet progeny at the posterior lip of the blastopore of the molluscs *Patella vulgata* (Lartillot et al., 2002a) and *Haliotis asinina* (Koop et al., 2007). Therefore, the *M. membranacea* data indicates that the molecular identity of the blastomeres remained conserved to a certain extent, despite the altered cleavage geometry and position of the second and third quartet in the bryozoan, placed more vegetally than spiral-cleaving embryos.

#### Fourth quartet: muscle and mesenchymal cells

The embryonic source of mesoderm in bryozoans remains a contentious topic (Gruhl, 2009). Classical works suggest that mesodermal cells derive from endodermal blastomeres, but could not demonstrate the embryonic origin with cellular resolution (Barrois, 1877; Calvet, 1900; Cor-rêa, 1948; d’Hondt, 1983; Pace, 1906; Prouho, 1892). However, recent ultrastructural data in *M.membranacea* suggests an ectodermal origin for the bryozoan mesoderm, from the delamination of an ectodermal cell during gastrulation (Gruhl, 2009). Our cell lineage data indicate that the first mesodermal cells of *M. membranacea* derive from the fourth quartet (4a–4c). The lateral cells 4a^A^ and 4c^A^ form the anterior muscles of the cyphonautes larva while the progeny of 4b^1^ gives rise to a stack of mesenchymal cells that express the anterior mesoderm marker *foxf*. We did not observe the delamination of an anterior ectodermal cell as described by Gruhl (2009), but cannot discard the existence of other cells contributing to the mesoderm of *M. membranacea*. Our work corroborates previous classical studies of bryozoan embryology by revealing that the mesoderm of the bryozoan *M. membranacea* originates from multiple fourth-quartet blastomeres.

The source of mesodermal tissues in spiral-cleaving embryos is extensively studied and discussed (Ackermann et al., 2005; Boyer et al., 1996; Boyer et al., 1998; Conklin, 1897; Gline et al., 2011; Hejnol, 2010; Hejnol et al., 2007; Henry and Martindale, 1998; Kozin et al., 2016; Lambert, 2008; Lartillot et al., 2002b; Lyons and Henry, 2014; Lyons et al., 2012; Lyons et al., 2015; Meyer et al., 2010; Render, 1997). There are generally two sources, an anterior mesoderm derived from the third quartet blastomeres (known as ectomesoderm) and a posterior mesoderm derived from the 4d blastomere. Even though most spiralians have the 4d as the sole endomesodermal contributor, there are exceptions – in the annelid *Capitella teleta* 3c and 3d generate mesoderm (Eisig, 1898; Meyer et al., 2010). The blastomeres contributing to the anterior mesoderm (usually 3a and 3b) are more variable (Hejnol et al., 2007; Lyons and Henry, 2014). This indicates that even within spiral-cleaving embryos, particular cell fates can shift to different blastomeres (Hejnol et al., 2007). We find that the source of mesodermal tissues of the bryozoan *M.membranacea* differs from other spiralians because (1) the third quartet does not contribute to the anterior mesoderm and (2) multiple blastomeres of the fourth quartet give rise to mesodermal tissues. In addition, the blastomeres 4a–4c often give rise to endodermal tissues in spiral-cleaving embryos (Hejnol et al., 2007; Henry and Martindale, 1998). Therefore, our data suggests that the specification of the anterior mesoderm in the bryozoan might have been shifted in time and in position from the third quartet to the fourth quartet.

Although only a subset of the genes we analyzed is expressed in the mesoderm of *M. mem-branacea* (*six3/6*, *evx*, *twist*, *foxf*), the patterns indicate that the bryozoan mesoderm is already regionalized at early gastrulation. For instance, we found that 4d expresses *evx* and *cdx*, genes commonly associated to posterior mesodermal and hindgut fates in spiralians (Fritsch et al., 2016; Fröbius and Seaver, 2006; Hiebert and Maslakova, 2015b; Hui et al., 2009; Kulakova et al., 2008; Le Gouar et al., 2003; Martín-Durán et al., 2015; Martín-Durán et al., 2016b; Rosa et al., 2005; Samadi and Steiner, 2010). Even though, we could not resolve the fate of the 4d blastomere in *M. membranacea*, the expression data indicates that the 4d blastomere might contribute to the posterior mesoderm and hindgut of the bryozoan. The lateral mesodermal cells 4a/4c and derivatives express *evx*, and possibly *twist*, in the same manner as the expression of *evx* orthologs in the annelid *Capitella teleta* during early development (Seaver et al., 2012). Finally, at the anterior mesoderm we find the expression of *foxf*, also observed in the brachiopod *Terebratalia transversa* (Passamaneck et al., 2015). Overall, these molecular data reveal that *M. membranacea* mesoderm is regionalized and that at least some of the expression patterns are conserved with other spiralians.

The 4d cell and its descendants also form the germline and are known to express *nanos* in spiral-cleaving embryos (Rebscher, 2014). Germ cells have not been identified during the embryogenesis of any bryozoan and were only found in zooids after metamorphosis (Reed, 1991). The expression of *nanos* in *M. membranacea* differs from the pattern of spiral-cleaving embryos, since *nanos* is expressed in two posterior cells of the second and third quartet. These blastomeres divide repeatedly, but *nanos* expression is always retained in two cells that become part of the larval internal sac – the structure that persists during metamorphosis giving rise to the outer case of the zooid (Stricker, 1988). Thus, we hypothesize the *nanos*-positive cells might be stem cells contributing to the differentiation of the internal sac, but further analysis in competent larvae and metamorphosed juveniles are needed to clarify the fate and molecular identity these cells.

#### 4Q blastomeres: gut

The 4Q blastomeres are the largest and yolkier cells of the gastrulating bryozoan embryo and in *M. membranacea*, they originate the cyphonautes larval gut. Similarly, the macromeres of spiral-cleaving embryos generally give rise to endodermal tissues (Ackermann et al., 2005; Child, 1900; Conklin, 1897; Hejnol et al., 2007; Henry and Martindale, 1998; Henry et al., 2004; Henry et al., 2010; Maslakova et al., 2004; Mead, 1897; Meyer et al., 2010; Wilson, 1892), even though in the polyclad *Hoploplana inquilina* these cells break up and the gut is likely derived from 4d (Boyer et al., 1998; Surface, 1907). Nevertheless the endodermal fate of these large vegetal cells appears to be a common feature of *M. membranacea* and spiral-cleaving embryos.

The expression of the endodermal marker *gata456b* corroborates the molecular identity of the 4Q blastomeres in the bryozoan. In spiralians *gata456* expression is mainly associated to endodermal and, in some cases, mesodermal tissues (Boyle and Seaver, 2008; Boyle and Seaver, 2010; Gillis et al., 2007; Martín-Durán et al., 2015; Martín-Durán et al., 2016b; Passamaneck et al., 2015; Wong and Arenas-Mena, 2016). The expression of *gata456b* in *M. membranacea* is clearly associated to the larval gut, suggesting that the molecular patterning of endodermal structures in the bryozoan is conserved with other spiralian groups, and possibly to other bilaterians (Martín-Durán and Hejnol, 2015). One exception is the bryozoan *Bugula neritina*, where *gata456* is expressed at the apical organ (Fuchs et al., 2011). However, this might be related to the fact that the coronate larva of *B. neritina* does not have a gut, and requires further investigation. The early expression of *gata456b* in *M. membranacea* does differ slightly from other spiralians, since neither the 4B blastomere nor the 4a–4c blastomeres show *gata456* transcripts (Boyle and Seaver, 2008; Boyle and Seaver, 2010; Wong and Arenas-Mena, 2016). Nevertheless, the common fate and *gata456* expression between *M. membranacea* 4Q blastomeres and the macromeres of other spiralians suggest that the bryozoan shares similar molecular and developmental traits for the endoderm patterning with spiral-cleaving embryos.

### A modified spiral cleavage

Our investigation shows a series of similarities between the embryonic development of the bryozoan *M. membranacea* and the embryogenesis of annelids, molluscs, nemerteans and polyclads. The vegetal blastomeres sequentially give rise to quartets of daughter cells, the first asynchronous cell divisions occur in the posterior quadrant, the quadrant identities can be identified at the 32-cell stage, the MAPK activity resembles that of equal-cleaving molluscs and several genes are expressed in equivalent blastomeres or embryonic regions (Figure 12). In addition, the early blastomeres of *M. membranacea* and of spiral-cleaving embryos have similar fates in the larval tissues (Figure 12). That is, the first animal blastomeres form the whole region from the apical organ to the corona – equivalent to the pretroch region – and the ciliated band itself, the second and third quartets contribute to the oral ectoderm, the fourth quartet gives rise to the mesoderm of the larva, and the four large vegetal blastomeres are internalized and become endoderm (Figure 12). Since the phylogeny of Spiralia indicates spiral cleavage is ancestral and bryozoans are nested within the clade (Kocot, 2016; Kocot et al., 2016; Laumer et al., 2015; Nesnidal et al., 2013; Struck et al., 2014), we interpret these developmental similarities as inherited traits from an ancestral spiral-cleaving embryogenesis.

In this context, during the evolution of gymnolaemate bryozoans the ancestral spiral cleavage pattern, characterized by the alternating oblique cell divisions, was modified to biradial cell divisions. While the cleavage pattern changed and the anterior mesoderm was reallocated to the fourth quartet, some aspects of the development have remained conserved, such as the D quadrant specification, MAPK activity and overall fate map of early blastomeres.

Spiral cleavage has been modified not only in bryozoans, but in different spiralian branches as well, such as flatworms, molluscs and brachiopods (Hejnol, 2010). In most of these cases, the embryonic development has changed to such extent that no traces of spiral cleavage are found, and the ancestral cleavage geometry can only be inferred by the phylogenetic position of the clade (e.g. the discoidal cleavage of cephalopods (Wadeson and Crawford, 2003)). Remnants of spiral cleavage usually consist of oblique mitotic spindles, as in the flatworm *Macrostomum lignano*, which displays a typical spiral cleavage pattern until the third cleavage, when the embryonic development becomes considerably modified (Willems et al., 2009). The bryozoan *M. membranacea* differs from these previously known cases because we can recognize shared cell lineage and developmental traits with spiral-cleaving embryos that are not the cleavage geometry itself. The evolutionary mechanisms involved in the transition from spiral to biradial cleavage remain unclear, but changes in the orientation of the mitotic spindle have a genetic basis and are beginning to be uncovered using molecular and computational approaches (Brun-Usan et al., 2016; Davison et al., 2016; Kuroda, 2015). Our data suggests that the quartet-divisions, cell fates and other traits commonly associated with a spiral cleavage program were maintained in the bryozoan development despite the evolutionary modification to a biradial cleavage pattern.

### Evolution of cleavage patterns

The fate of the early embryonic blastomeres is thought to be causally associated to the cleavage pattern during development (Valentine, 1997; Wray, 1994). In such case, a change in the cleavage geometry would lead to a change in the cell fates. However, the bryozoan cell lineage illustrates a case where the cleavage pattern and blastomere fates are not evolutionary coupled. We find that the fates of the early blastomeres are similar between *M. membranacea* and spiral-cleaving embryos despite the modified biradial cleavage pattern. The relative positioning of the second and third quartets even differs between the bryozoan and a typical spiral-cleaving embryo (Figure 12), but these blastomeres still contribute to similar tissues, suggesting the early cell fate determination remained relatively conserved during bryozoan evolution. The bryozoan cell lineage illustrates how a widely conserved determinate cleavage pattern – spiral cleavage – can evolve without major changes in other developmental traits, such as the blastomere fates and molecular identity.

We found several developmental genes expressed in a similar spatial arrangement between bryozoans and other spiralians, as revealed by MAPK (3D blastomere), *otx* (basal blastomeres of the first quartet), and *bra* and *foxa* (second and third quartets). This molecular map is similar not only to the typical spiral-cleaving embryos, but also to brachiopod embryos (Martín-Durán et al., 2016b; Passamaneck et al., 2015), whose embryos have a much greater number of cells and no stereotypic cleavage pattern (Long and Stricker, 1991). Thus, a single cell in the bryozoan embryo expressing *gata456* might be homologous to a whole region of *gata456* expression in the brachiopod embryo (Passamaneck et al., 2015), as suggested by Hejnol (2010). This reinforces the hypothesis that cell fate determination is not tied to a particular cleavage pattern, but depends on the underlying molecular framework established early in development (Henry et al., 1992). This is the case for nematodes, whose cleavage patterns diverged drastically between groups without a corresponding change in the resulting phenotype (Schulze and Schierenberg, 2011).

However, a clearer parallel case to the spiral-to-biradial evolution is the transition from the spiral cleavage of polychaete annelids to the derived cleavage of clitellate annelids; despite the differences in the cleavage pattern, the clitellate fate map does not deviate significantly from the annelid ground plan (Kuo, 2017). Overall, our findings support the hypothesis that in evolutionary terms, the causal ontogenetic connection between cleavage pattern and blastomere fates, if any, can be broken (Scholtz, 2005). In the case of the bryozoan *M. membranacea*, the molecular identity and fate of the early blastomeres might have been maintained, despite the modification in the geometry of cell divisions. Further comparative cell lineage studies with other non-spiral spiralian lineages, such as gastrotrichs and rotifers, will be crucial to establish the ancestral traits of spiralian development and to better comprehend the relation between cleavage patterns and cell fates during evolution.

## Conclusions

The embryonic development of *M. membranacea* provides a unique comparative standpoint to the typical spiral cleavage pattern. It reveals that spiral cleavage is not an all-or-nothing character and has been extensively modified in the diverse spiralian lineages. In particular, we suggest that the cleavage geometry of the bryozoan embryo evolved independent from other spiralian developmental traits. Therefore, modifying spiral cleavage does not require drastic developmental changes such as the ones found in cephalopods or parasitic flatworms. More generally, our data suggests that determinate cleavage patterns can be modified without major changes in the identity of blastomeres and cell fates, which alleviates the idea that the cleavage pattern is evolutionary coupled to the specification of cell fates. In this perspective, the evolutionary conservation of cell fates in spiral-cleaving clades might be a consequence of a conserved underlying molecular patterning, overlaid by a determinate cleavage pattern. Overall, our work highlights the importance of comparative data to better understand the evolution of spiralian development.

## Methods

### Collection, spawning and cultures

We collected *M. membranacea* in the fjord waters of Hjellestadosen (60°15'23.9" N 5°14'20.1" E) in Bergen, Norway, between May and September. We handpicked kelp blades with ripe bryozoan colonies from floating boat docks, and maintained the leaves in tanks with flowing sea water at 10 °C. To induce spawning, we cut a portion of the kelp blade with mature colonies, usually the ones with more opaque whitish/pinkish zooids, and transferred it to glass bowl with sea water sterilized with UV-light and filtered through a 0.2 µm mesh (UVFSW). The bowl was placed under a stereomiscroscope with direct light and a digital thermometer to monitor the water temperature. Ripe colonies begin to spawn in around five minutes or more, or usually when the temperature reached 15 °C. Once the temperature rose to 16 °C, the bowl was cooled down on ice with no direct light. The spawning colony was sequentially transferred to new bowls with UVFSW at 10 °C to distribute the vast amounts of eggs. For each bowl with eggs, we added EDTA to a final volume of 0.1mM (usually ~20 µL of 0.5M EDTA) to induce egg activation (Reed, 1987). The bowl was then placed in a incubator at 15 °C for 30–60 min. Activated eggs were concentrated by swirling, distributed to smaller glass bowls with UVFSW and washed twice to remove the EDTA. We adjusted the amount of eggs per bowl to avoid that the eggs sitting on the bottom of the dish touch each other. We kept the cultures at 15 °C. One colony could be re-used for spawning multiple times. *M. membranacea* colonies maintained in the flowing tanks remained viable to developmental studies for a week.

### 4D recordings and cell tracing

We pipetted embryos at the 2-cell stage to a glass slide coated with poly-l-lysine. We mounted the embryos under a cover slip with supporting clay feet, completed the volume with UVFSW and sealed the cover slip with vaseline. The slide was put under an automated 4D microscopy system (Caenotec) (Hejnol and Schnabel, 2006) with a cooling ring around the objective to keep the temperature at 15 °C. We recorded full-embryo stacks (40–60 optical slices) every 40s under differential interference contrast. Development was recorded for approximately 24h, when the embryo became ciliated and swam away from the field of view.

Raw data consists of a sequence focal levels for each time point. We loaded the data into the tracking software SIMI°BIOCELL (SIMI®) and manually traced individual cells. The results of this manuscript are compiled from the cell tracking data of four different embryos (Figure S8). Source files with the cell lineages and blastomere synchrony analyses are available at Vellutini et al. (2017).

### Cell lineage nomenclature

To analyze and compare the cell lineage of *M. membranacea* we annotated the individual cells with a modified spiral cleavage nomenclature (Child, 1900; Conklin, 1897; Wilson, 1892). The identical blastomeres at the 4-cell stage were labeled as A, B, C and D according to their correspondent fate, as identified by the video recordings. In such case, the D quadrant was assumed to be the blastomere giving origin to the dorsal/posterior region of the cyphonautes larva. At the 8-cell stage, the animal blastomeres were named 1a–1d and the vegetal blastomeres 1A–1D. The first cleavage of these animals blastomeres give rise to the first octet, referred with the subscript _o_ (e.g. 1qo refers to all cells of the first octet). An octet is made from four inner cells, indicated by the subscript _i_ (for internal), and four outer cells indicated by the subscript _e_ (for external). Derivatives of the animal octets received the standard superscripts of spiral cleavage with ^1^ for the apical progeny and ^2^ for the basal progeny (e.g. 1a_i_^1^ refers to the inner A quadrant cell of the apical progeny from the first octet). The quartets derived from the four large blastomeres were labeled as in spiral cleavage (2q–4q). And the progeny of the first twelve-tet followed the spiral cleavage^1^ and ^2^ superscripts. Finally, we labeled meridional with the superscript ^R^ for the cell to the right and ^L^ for the cell to the left, when viewed from the egg axis.

### Fixation methods

We fixed representative developmental stages for antibody staining in 4% formaldehyde for 1h at room temperature, washed the embryos in PTw (1x PBS + 0.1% Tween-20) and stored (in PTw) at 4 °C. For in situ hybridization, we fixed the samples in a solution of 4% formaldehyde / 0.2% glutaraldehyde solution to avoid tissue damage during the protocol. After 1h fixation at room temperature, we washed the embryos in PTw, dehydrated through methanol series and kept the samples in 100% methanol at −20 °C.

### Gene cloning and in situ hybridization

We assembled the Illumina reads from *M. membranacea* (SRX1121923) with Trinity (Grabherr et al., 2011) and used known genes to identify putative orthologs in the transcriptome. We performed PCR gene specific primer pairs on cDNA synthesized with the SMARTer RACE cDNA Amplification kit (Clontech). Primers were designed with Primer3 (Untergasser et al., 2012). We synthesized antisense DIG-labeled riboprobes with MEGAscript kit (Ambion) and performed colorimetric *in situ* hybridization according to an established protocol (Martín-Durán et al., 2012). Sequences were deposited in the GenBank (NCBI) with the accession numbers XXXXXXX-XXXXXXX.

### Gene orthology

Orthology was assigned by aligning amino acid sequences of *M. membranacea* against annotated genes from diverse metazoans using MAFFT 7.271 (Katoh and Standley, 2013), retaining only informative portions of the alignment with GBlocks 0.91b with relaxed parameters (Talavera and Castresana, 2007) and running a Maximum Likelihood phylogenetic analysis with RAxML 8.2.4 (Stamatakis, 2014) using automatic model recognition and rapid bootstrap. Alignments were manually verified using UGENE (Okonechnikov et al., 2012). Resulting trees from the maximum likelihood analysis were rendered into cladograms using the ETE Toolkit (Huerta-Cepas et al., 2010) (Figure S9). Gene orthology runs and source files are available at Vellutini et al. (2017).

### Immunohistochemistry and MAPK antibody

We permeabilized the embryos with several washes in PTx (1x PBS + 0.2% Triton X-100) for 2h and blocked with two washes of 1h in PTx + 0.1% BSA (Bovine Serum Albumin) succeeded by 1h incubation in PTx + 5% NGS (Normal Goat Serum). Samples were incubated with the primary antibody for the MAPK diphosphorylated ERK-1&2 (Sigma M9692-200UL) diluted 1:200, and stored overnight at 4 °C on a nutator. We removed the MAPK antibody with three 5 min and four 30 min washes in PTx + 0.1% BSA, blocked in PTx + 5% NGS for 1h and incubated nutating overnight at 4 °C with the secondary antibody Anti-Mouse-POD conjugate (Jackson) diluted 1:250. We removed the secondary antibody with three 5 min followed by several washes in PTx + 0.1% BSA for 2h. To amplify and develop the signal, we incubated the embryos for 3–5 min with the provided reagent solution and fluorochrome from TSA reagent kit Cy5 (Perkin Elmer). We stopped the reaction with two washes in a detergent solution (50% formamide, 2x SSC, 1% SDS) at 60 °C, to reduce background, followed by PTw washes at room temperature. We stained nuclei by incubating permeabilized embryos in DAPI 1:500 or Sytox Green 1:1000 for 2h. Nuclei staining was combined with f-actin staining by the addition of BODIPY FL Phallacidin 5 U/mL previously evaporated to remove methanol.

### Microscopy and image processing

We mounted in situ embryos in 70% glycerol in PTw. Embryos from antibody staining were mounted in 97% 2,2’-Thiodiethanol (Asadulina et al., 2012; Staudt et al., 2007), 80% Glycerol in PBS or SlowFade^®^ Gold Antifade (ThermoFisher). We imaged the samples with a Zeiss Axio-Cam HRc mounted on a Zeiss Axioscope A1, using differential interference contrast technique (Nomarski) for in situ hybridizations and a fluorescent lamp for the MAPK antibody staining. We used a Confocal Leica TCS SP5 to image fluorescent samples. Colorimetric in situ hybridizations were also scanned under the confocal using reflection microscopy (Jékely and Arendt, 2007). We processed all resulting confocal stacks in Fiji (Schindelin et al., 2012). When necessary, we adjusted the distribution of intensity levels to improve contrast with Fiji or GIMP. We created vector graphics and assembled the figure plates using Inkscape.

## Acknowledgements

We thank the S9 members for fruitful feedback and discussions, in particular Anette Elde, Jonas Bengtsen and Anlaug Boddington for helping with the bryozoan collections, and Sabrina Schiemann for the help with the 4D microscopy. The study was funded by the core budget of the Sars Centre and by The European Research Council Community’s Framework Program Horizon 2020 (2014–2020) ERC grant agreement 648861 to AH.

### Competing interests

We declare to have no competing interests.

## Figures

**Figure S1.**
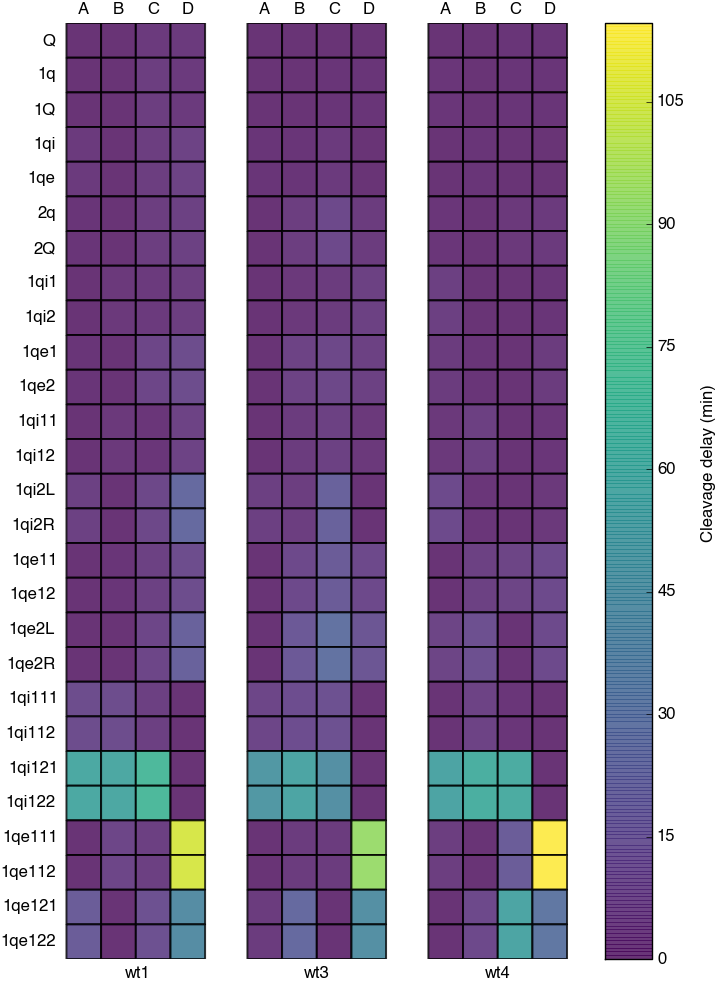
Synchrony in the cleavage of correspondent quartets in three *M. membranacea* embryos. The color gradient represents the time a cell took to divide after the first cell of its quartet had divided. Each column represents one embryo that has been tracked from the animal pole (wt1, wt3 and wt4). The most significant and consistent asynchronous events occur in the quartets 1q_i_^12^ and 1q_e_^11^.

**Figure S2.**
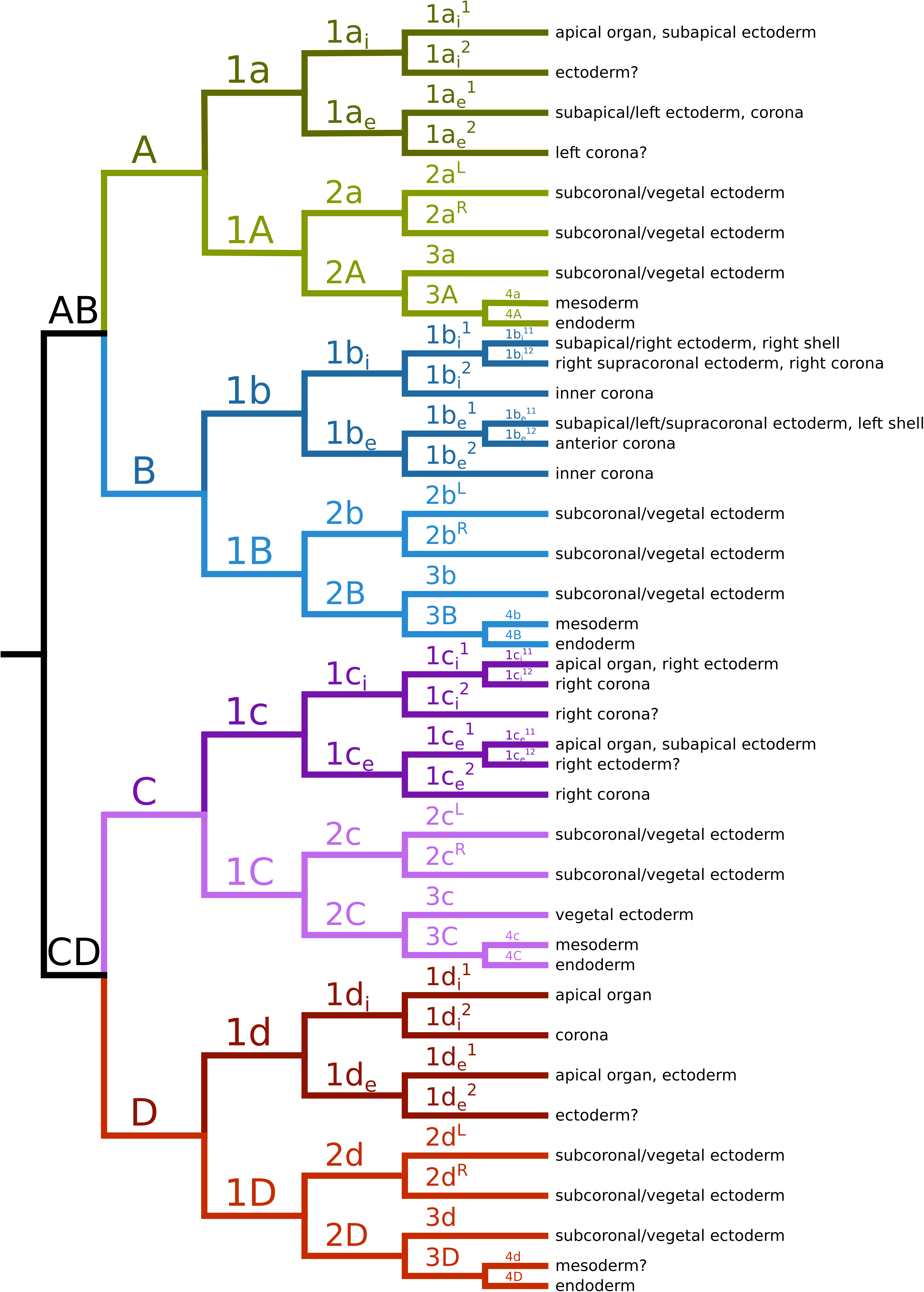
Detailed fate map of *M. membranacea*.

**Figure S3.**
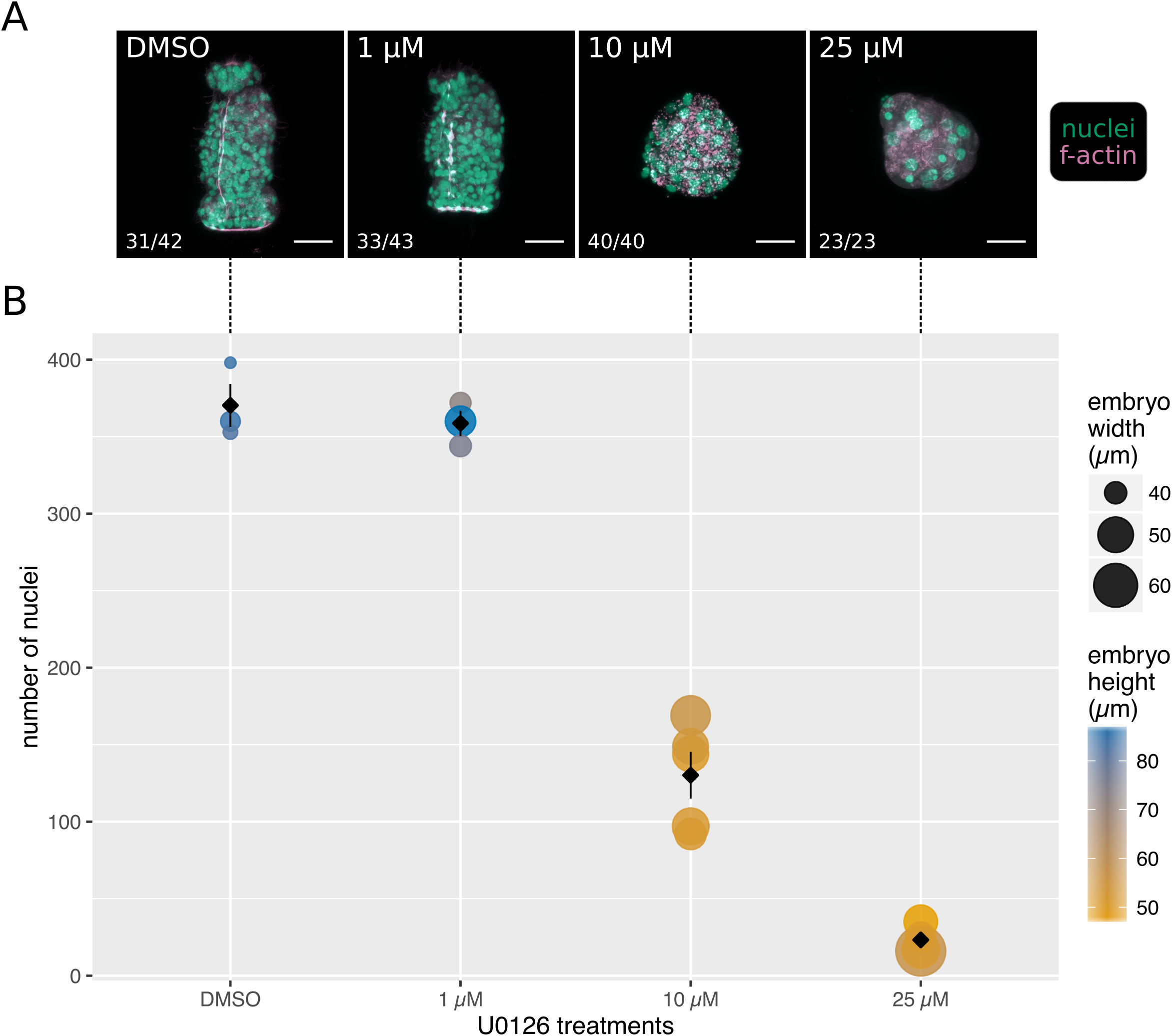
Development of *M. membranacea* under different concentrations of the MEK inhibitor U0126. (A) Maximum intensity projection of a confocal stack for the most representative phenotype of each U0126 treatment. Ratio in the lower left corner shows the number of embryos scored for the shown phenotype versus the total number of embryos in the treatment. Phenotypes in an additional seawater-only treatment (no DMSO) were indistinguishable from DMSO control and had ratio of 31/37 (not shown). (B) Measurements for the number of nuclei (y axis), embryo width (point size) and embryo height (color scale) for the confocal scans of (A). Each colored point represents one embryo, the black rhombus and error bars shows the mean number of nuclei with standard error. Number of embryos scanned and measured per treatment: DMSO=3, 1 µM=3, 10 µM=5, 25 µM=4. In all treatments, U0126 was added at 3 hpa (2-cell stage), the embryos developed at 10 °C and were fixed at 44 hpa. Scale bars = 20 µm.

**Figure S4.**
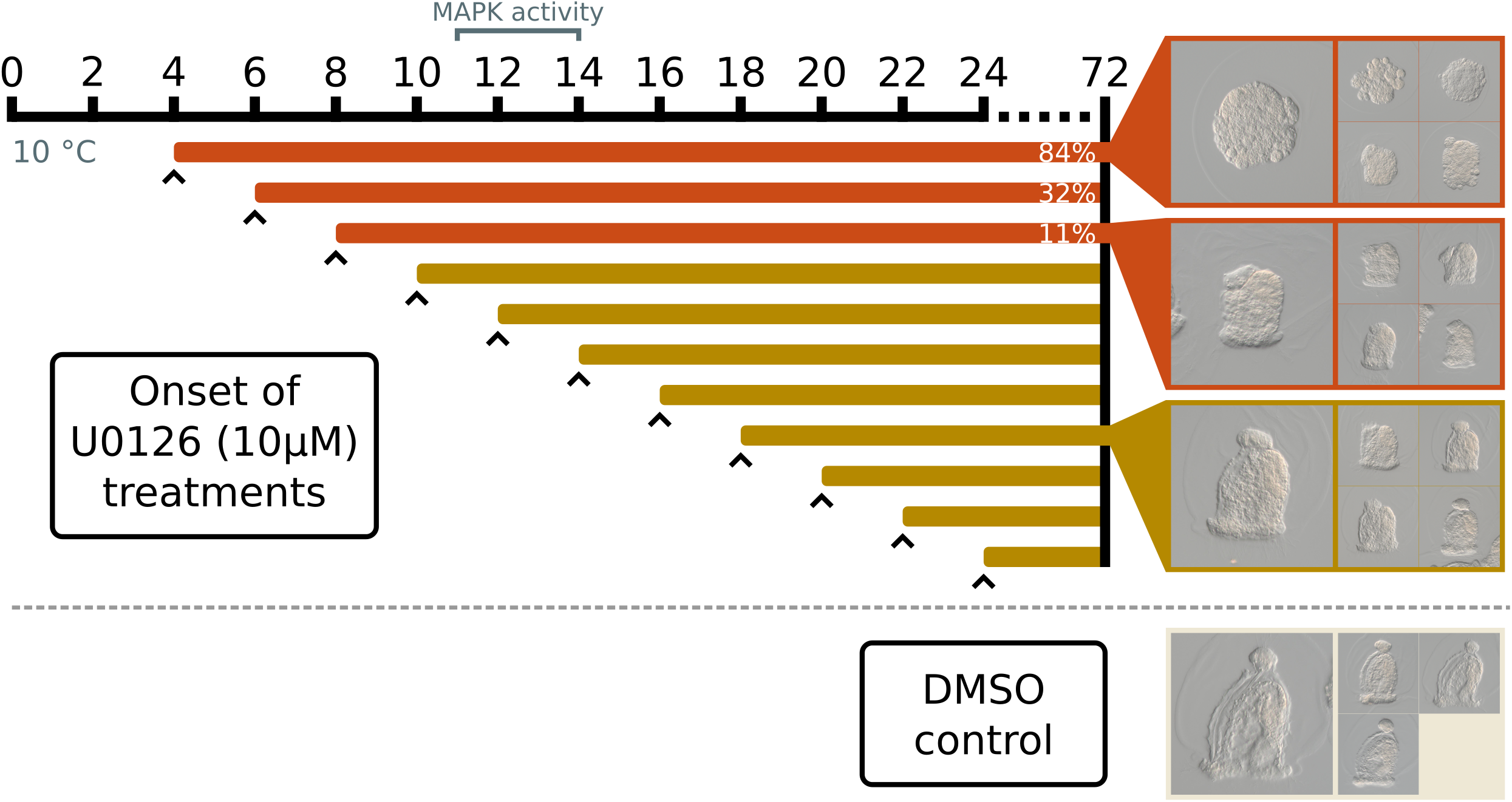
*M. membranacea* embryos treated with the MEK inhibitor U0126 (10 µM) from different developmental stages. All treatments developed at 10 °C and were fixed at 72 hpa. Arrowhead indicates when the U0126 treatments began for each experimental condition, represented by the horizontal colored lines. Representative phenotypes are shown for the 4, 8 and 18 hpa treatments. We scored 100 embryos under light microscopy for each treatment to obtain the ratio of severe/mild phenotypes. Treatments showing the severe phenotype are shown in orange, with the percentage of severe phenotypes indicated at the right end. Treatments without severe phenotypes were colored in yellow.

**Figure S5.**
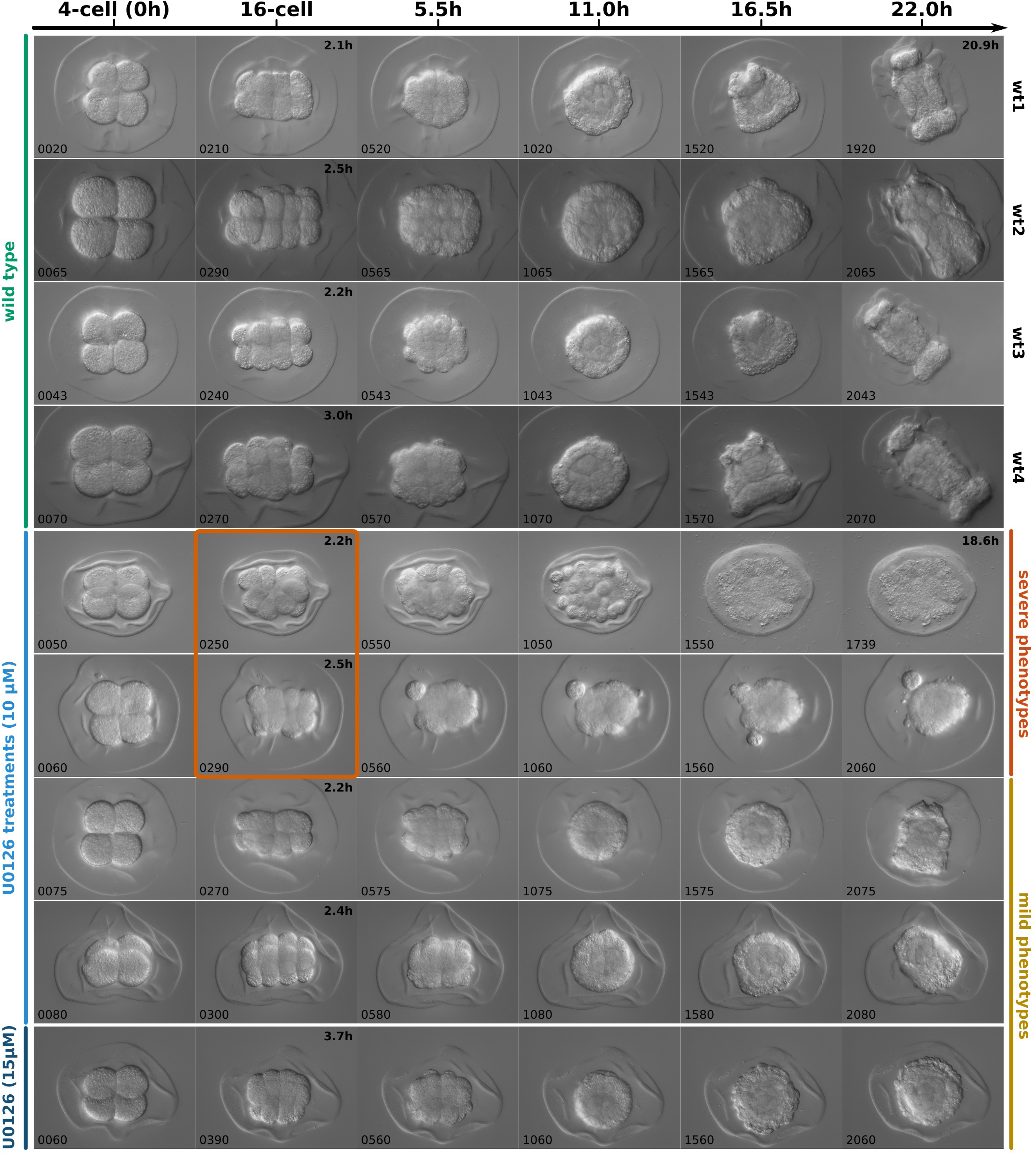
4D recordings of *M. membranacea* embryos treated with the MEK inhibitor U0126.Each row corresponds to the timeline of an individual embryo. All recordings were synchronized by the timing of the 2nd cleavage (4-cell stage = 0h). Time scale shows the number of hours after 4-cell stage. The exact developmental time is shown on the top right corner of panels that do not correspond to the time shown in the main scale. Frame number of each panel is shown on the bottom left corner (a frame was captured every 40 s). The orange rectangle indicates the cleavage abnormality observed in embryos that exhibit the severe phenotype.

**Figure S6.**
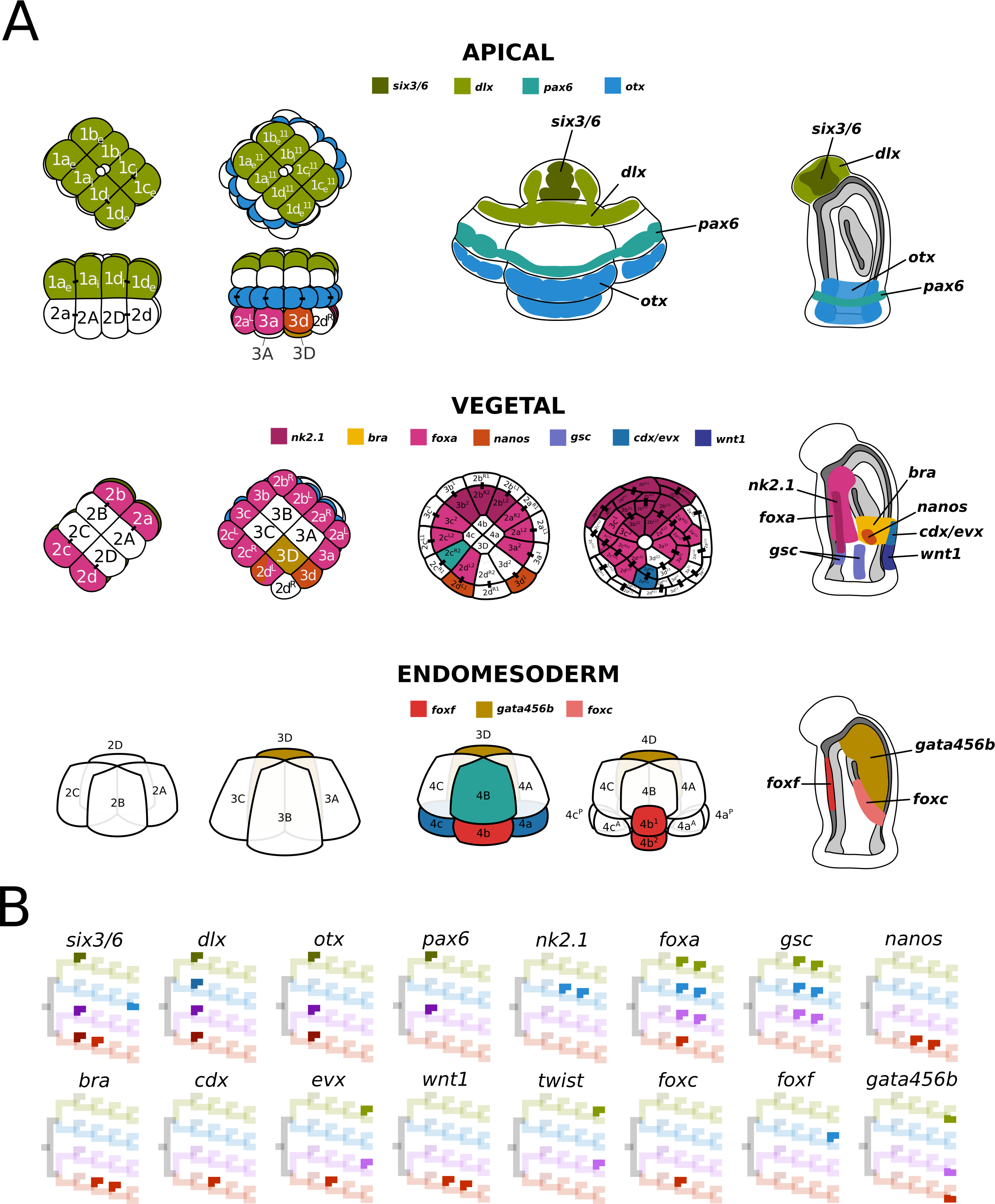
Gene expression throughout the cell lineage of *M. membranacea*. (A) Various developmental stages illustrating with cellular resolution, whenever possible, the expression of candidate genes in the animal ectoderm, vegetal ectoderm and endomesoderm. (B) Cell lineage diagrams indicating in which lineages the genes are being expressed. Gene activity is indicated in the branches with vivid colors.

**Figure S7.**
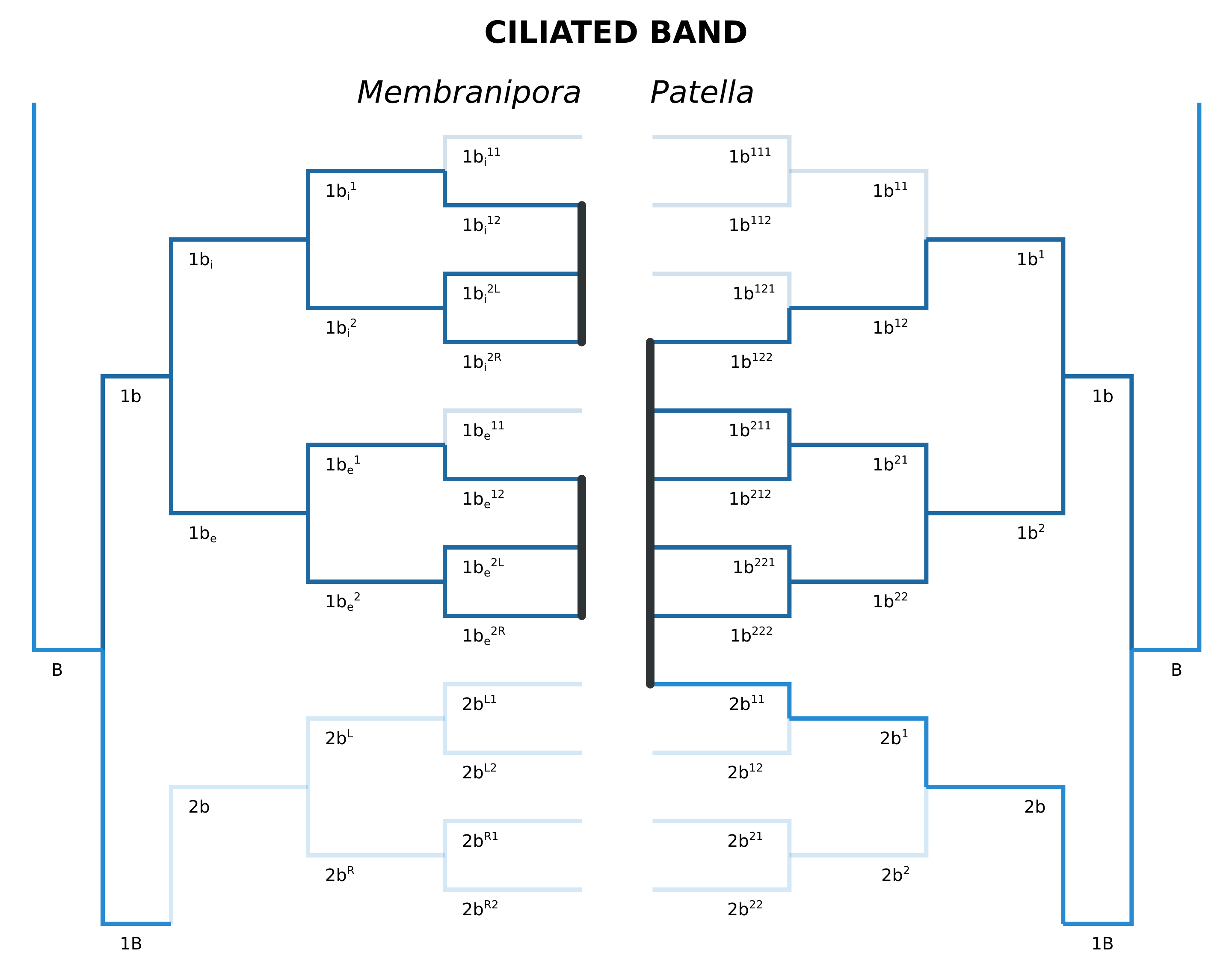
Cell lineage comparison between the larval ciliated bands of *M. membranacea* and *Patella vulgata* (Damen and Dictus, 1994).

**Figure S8.**
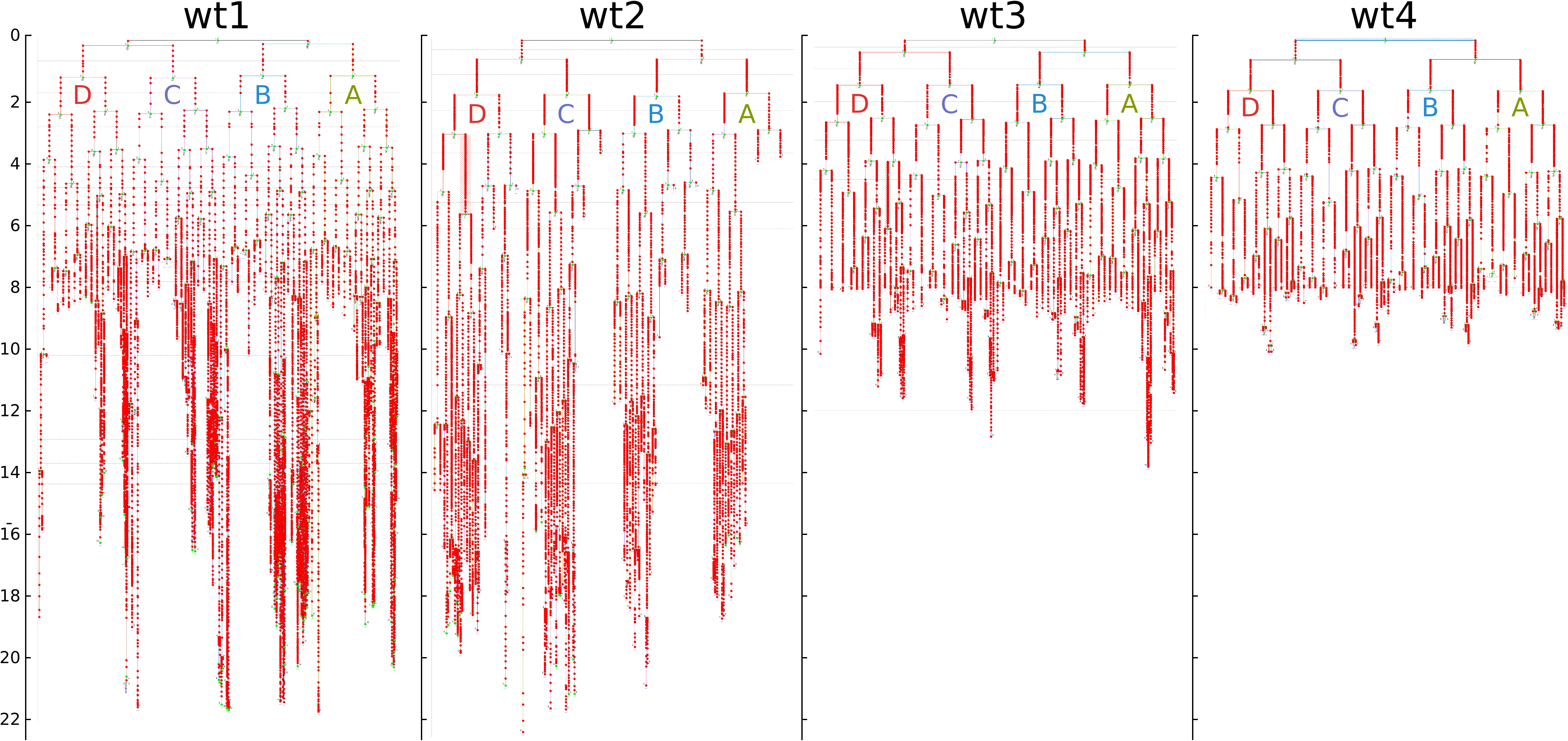
Cell lineages for three wild type (wt) embryos of *M. membranacea*. **wt1** is a recording of the animal pole providing most of the data for the aboral epithelium, **wt2** is a vegetal pole view providing detailed information for the vegetal ectoderm, and **wt3** and **wt4** are additional recordings of the animal pole. Development time measured in hours post activation (hpa) is shown in the Y axis.

**Figure S9.**
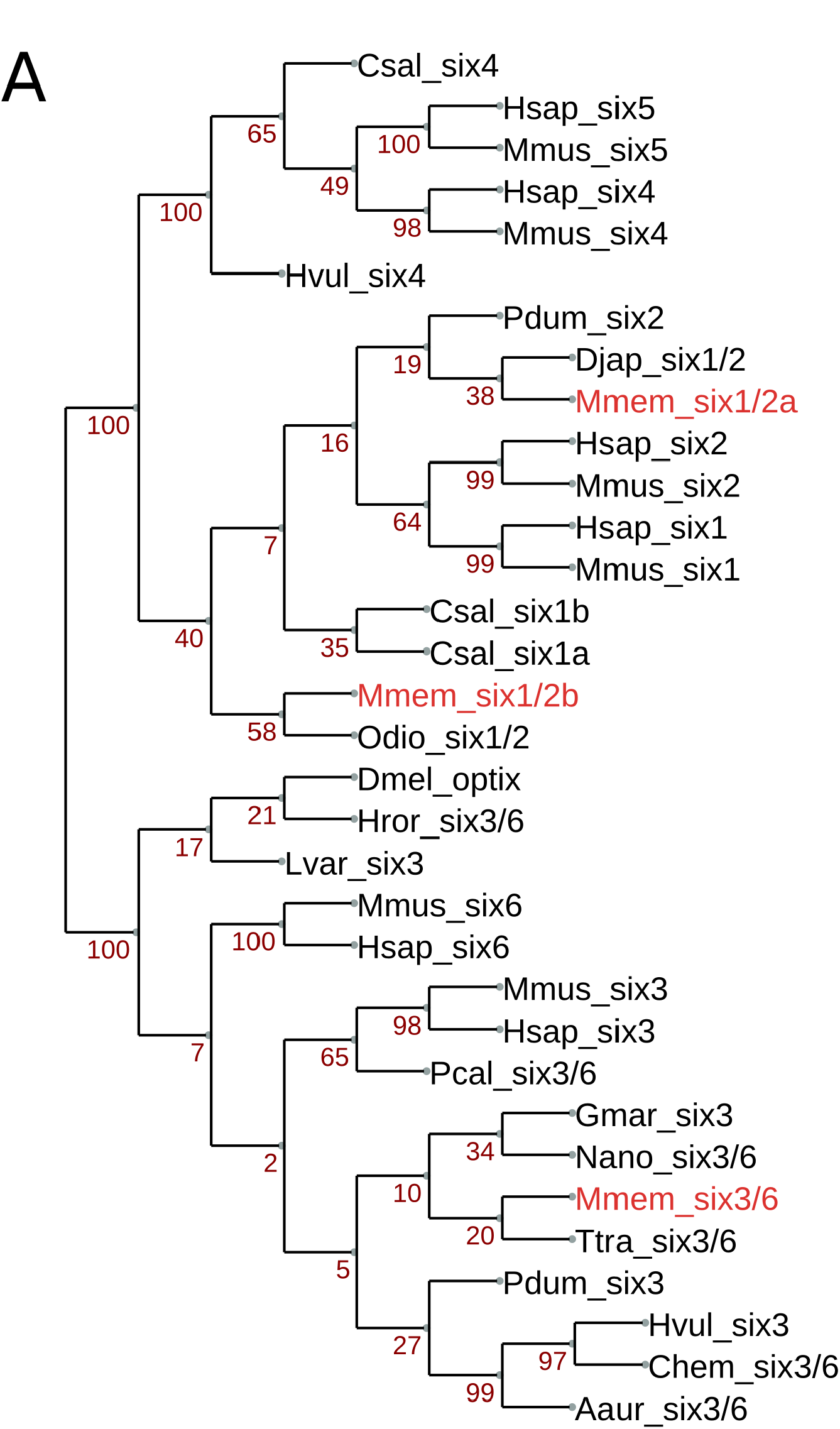
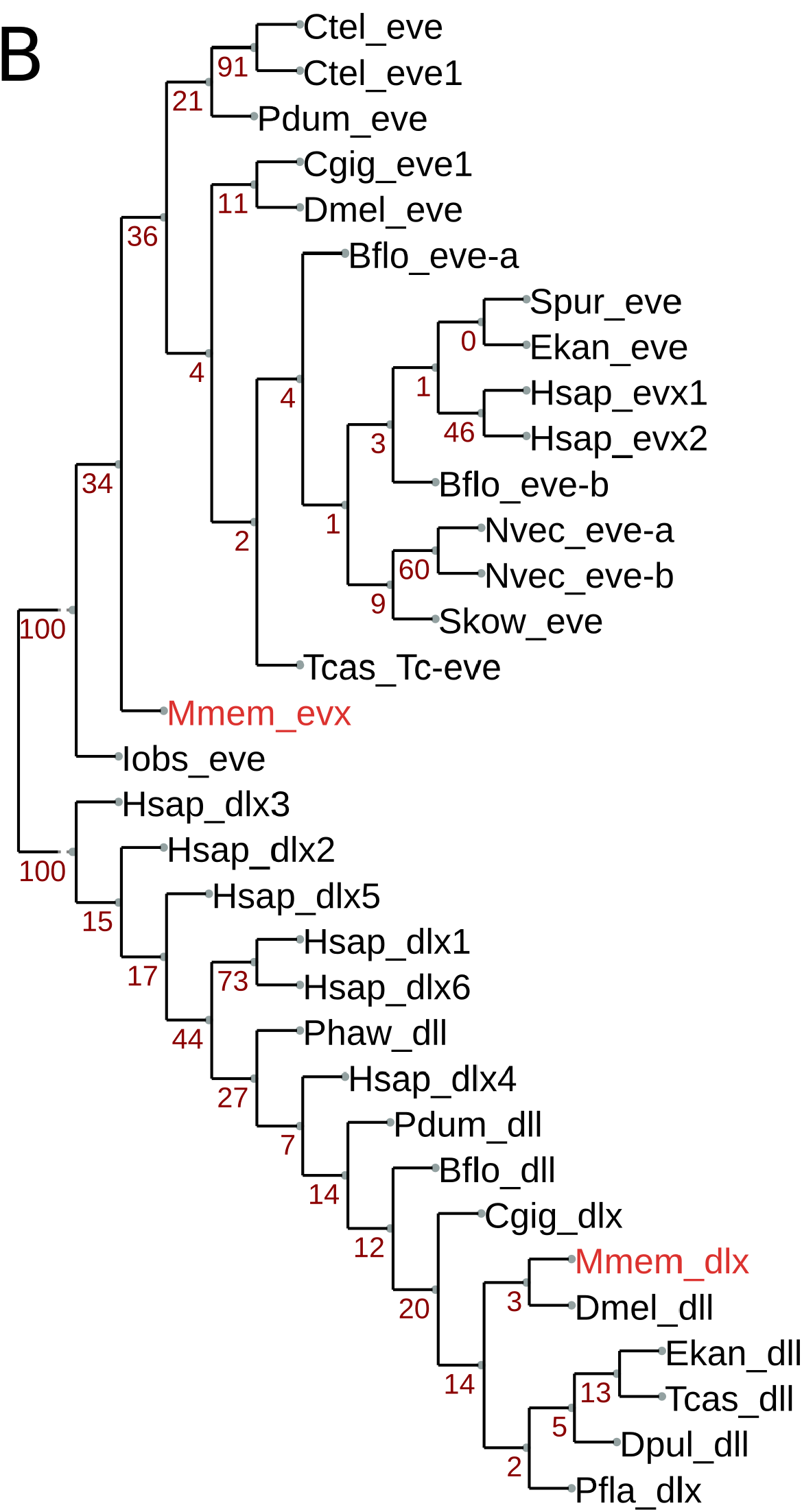
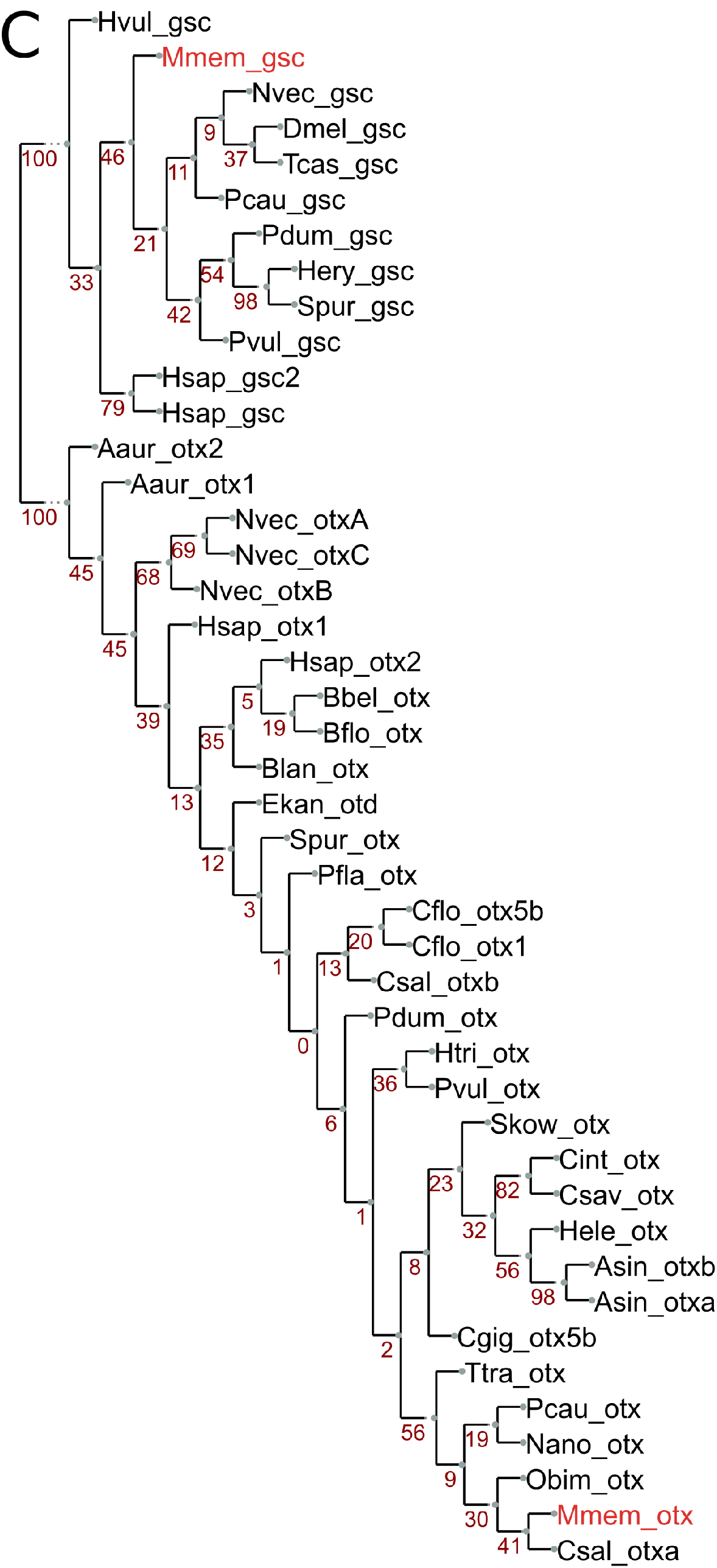
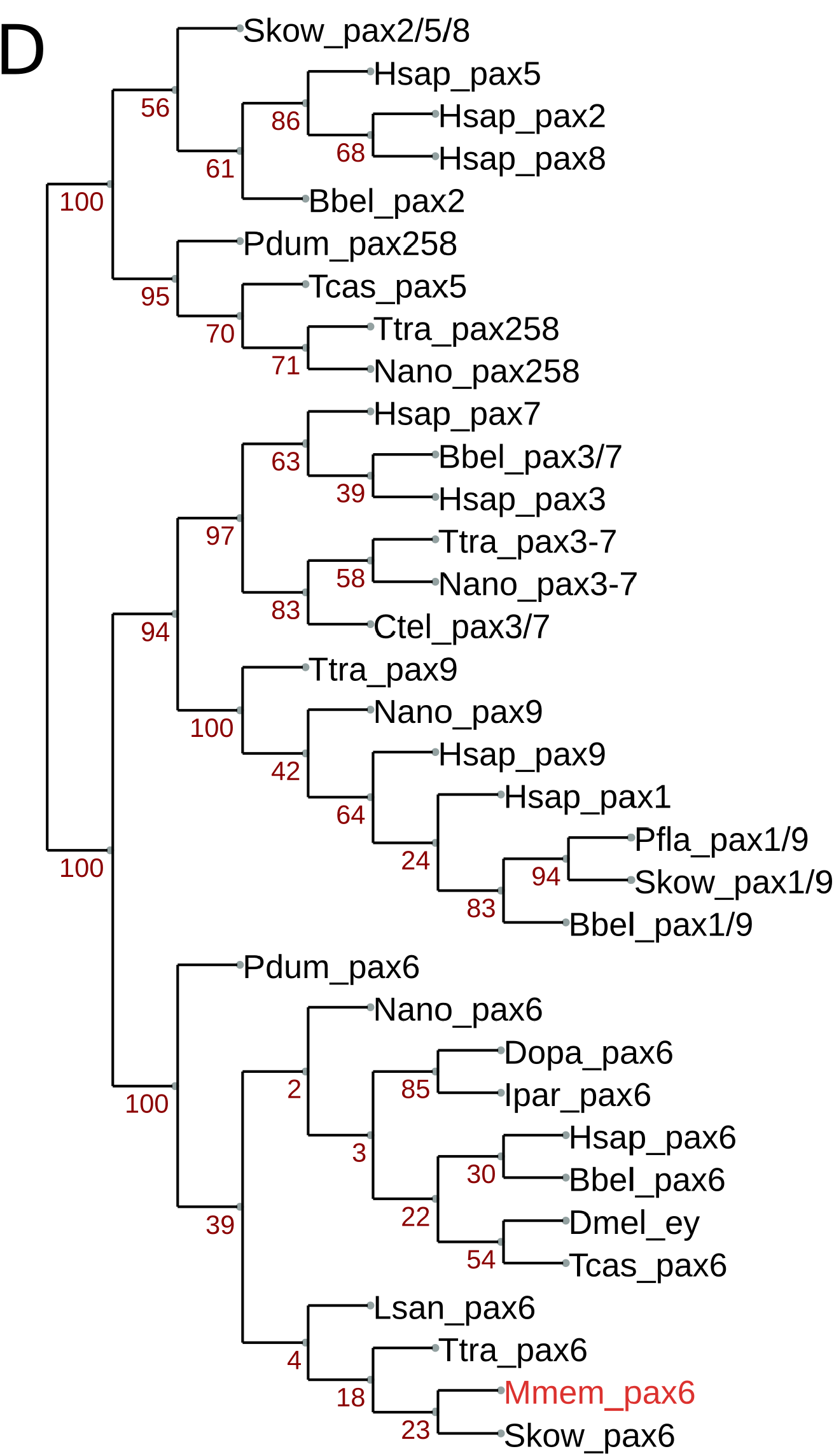
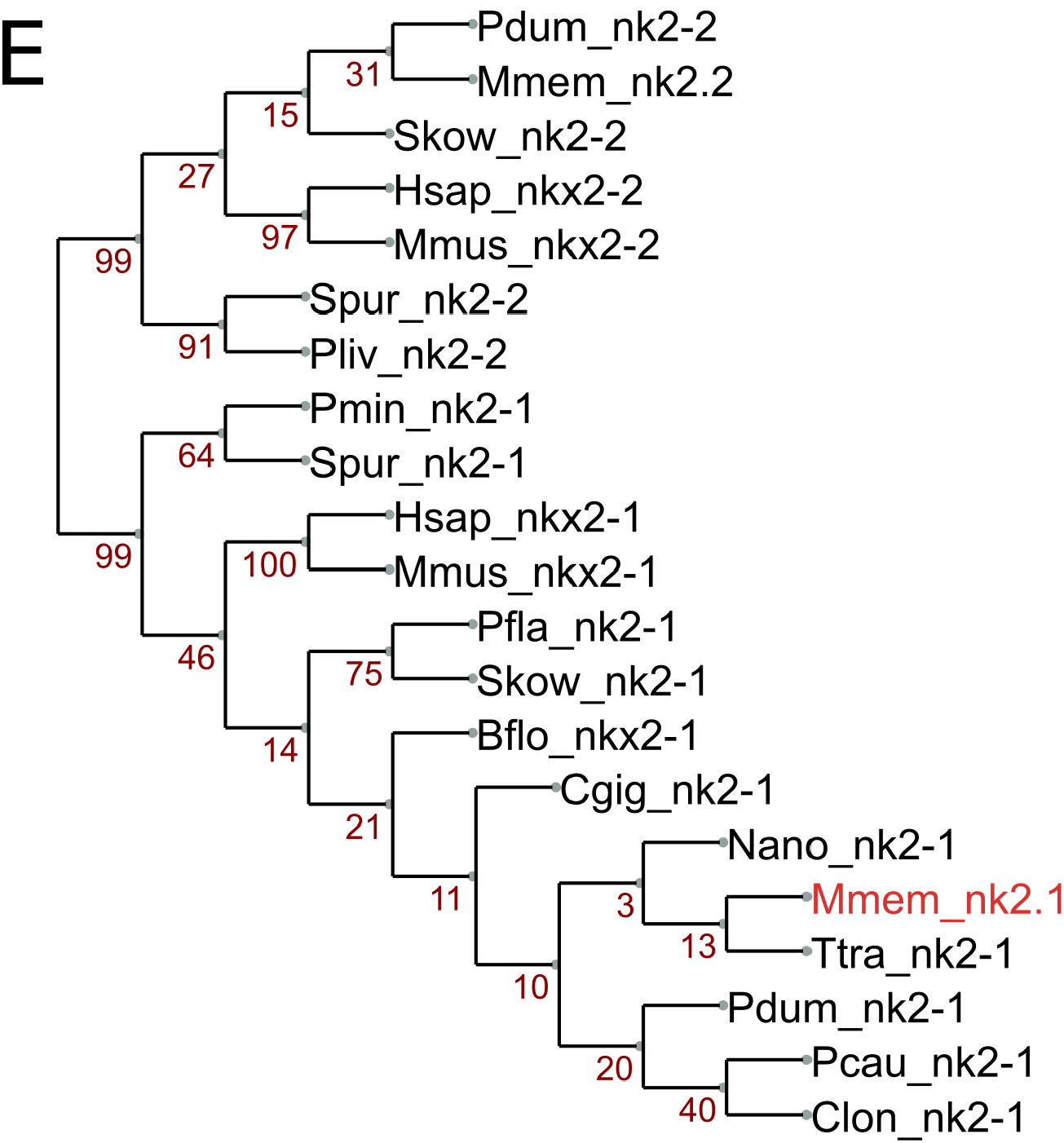
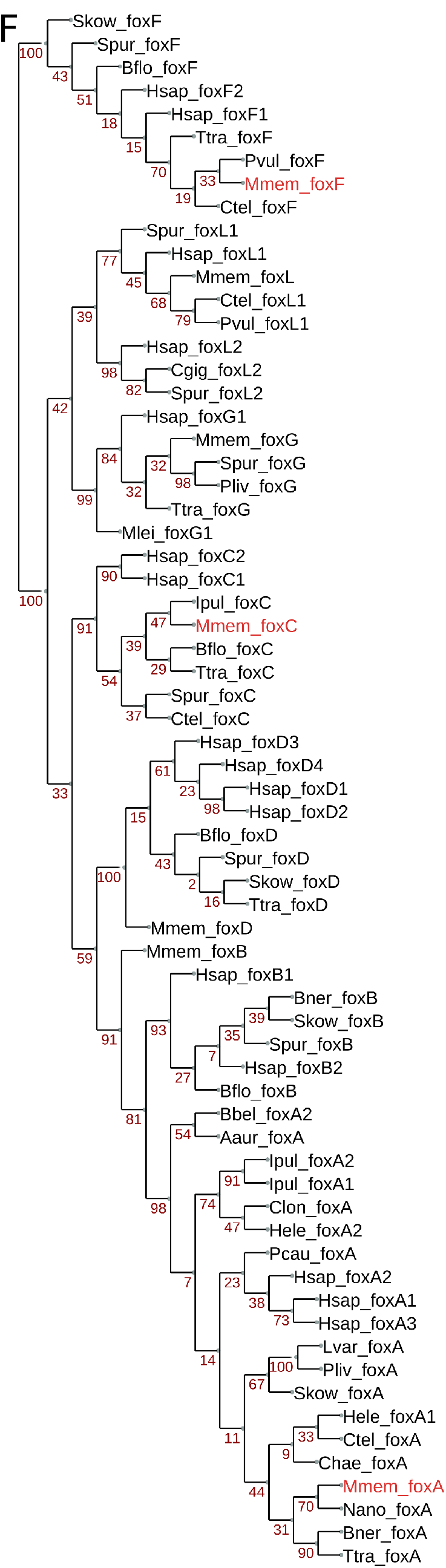
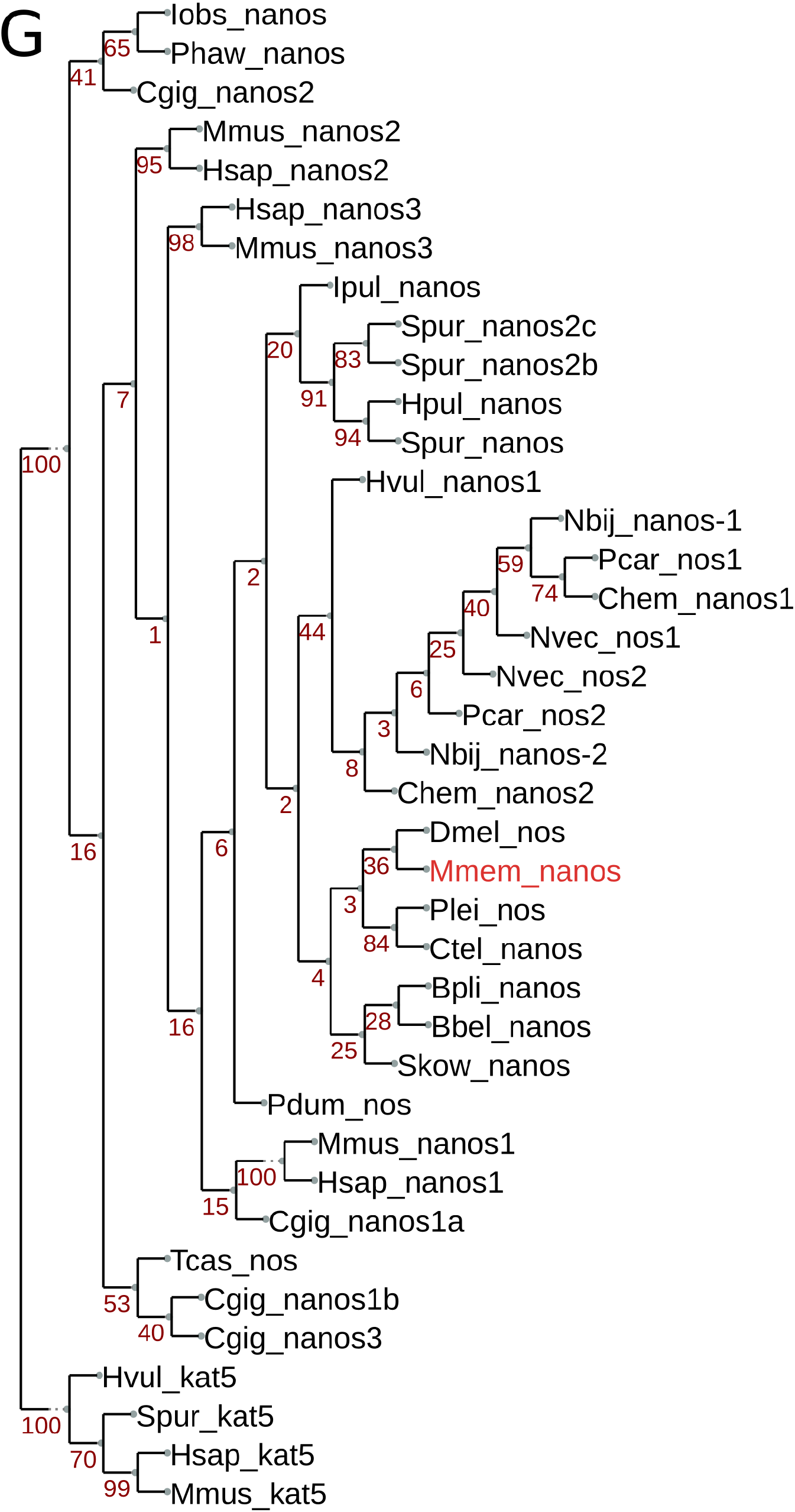
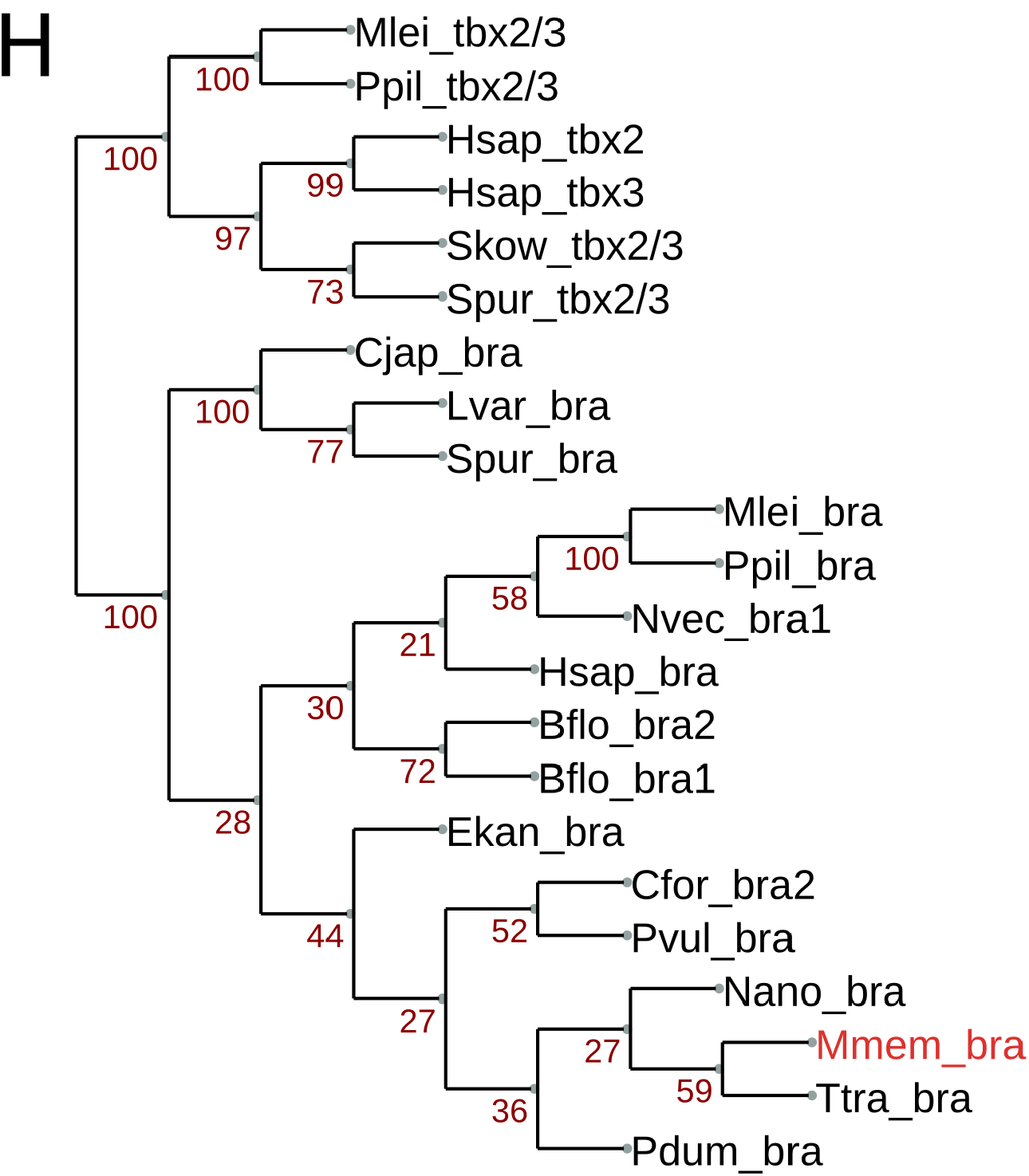
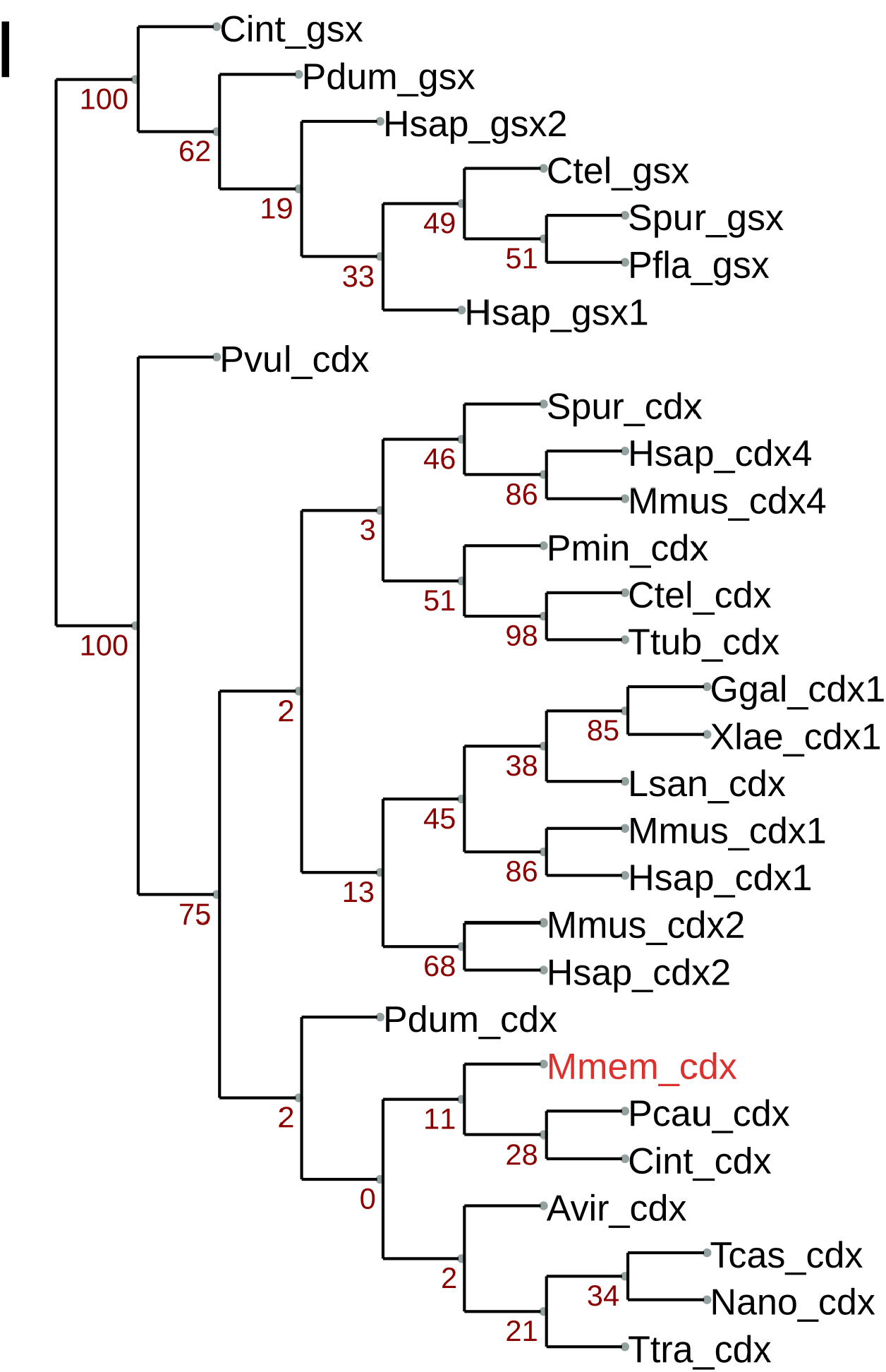
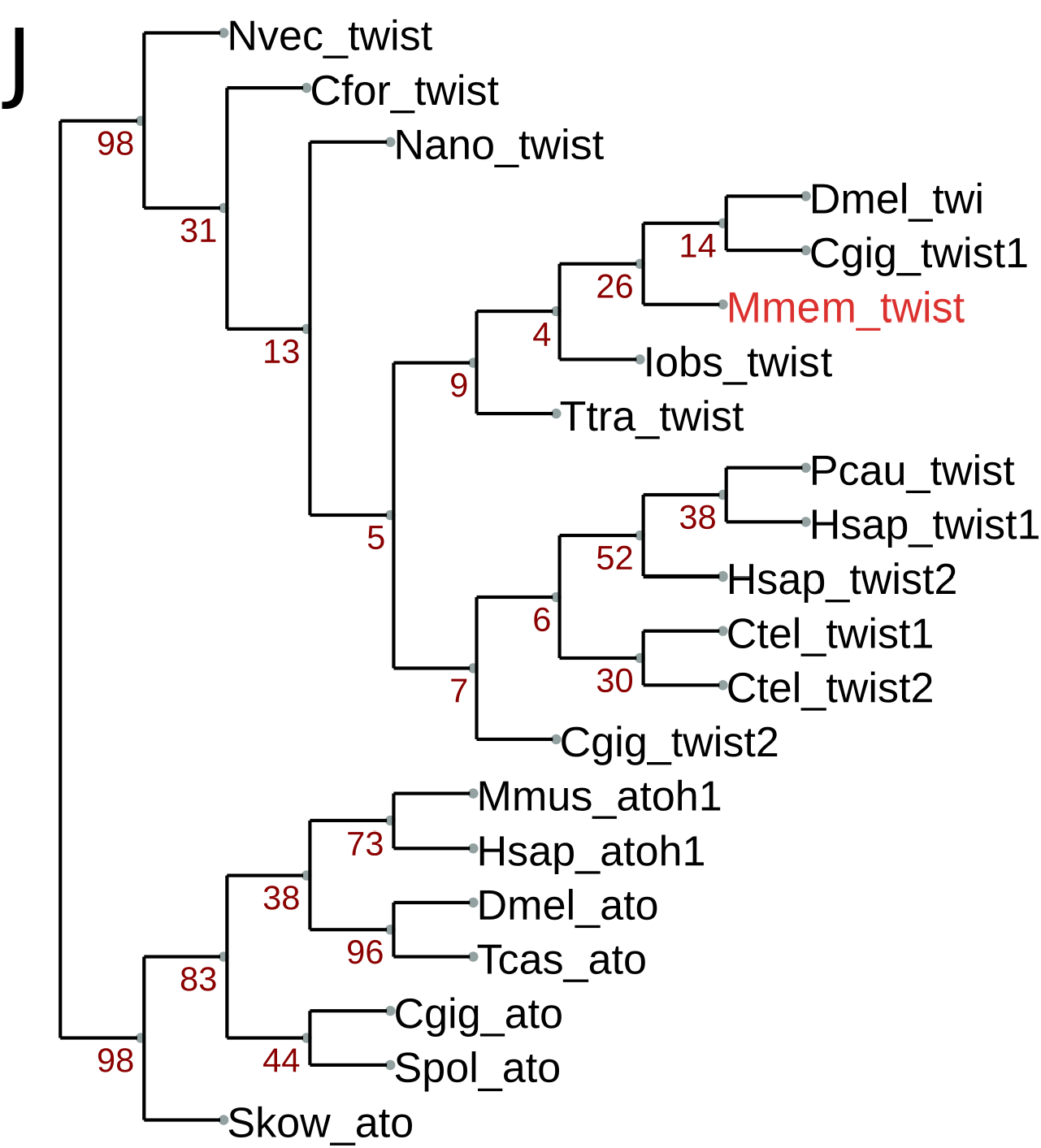
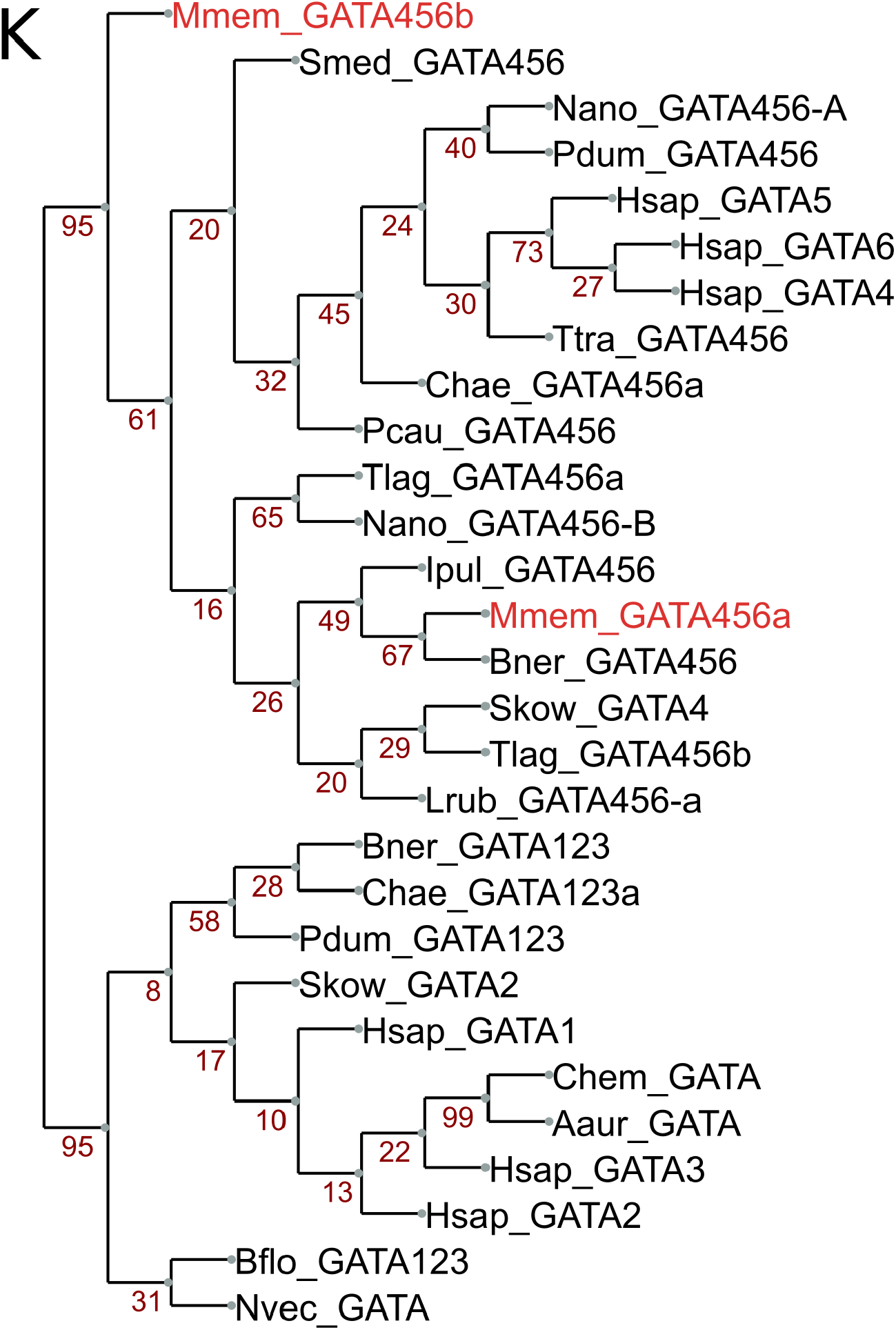
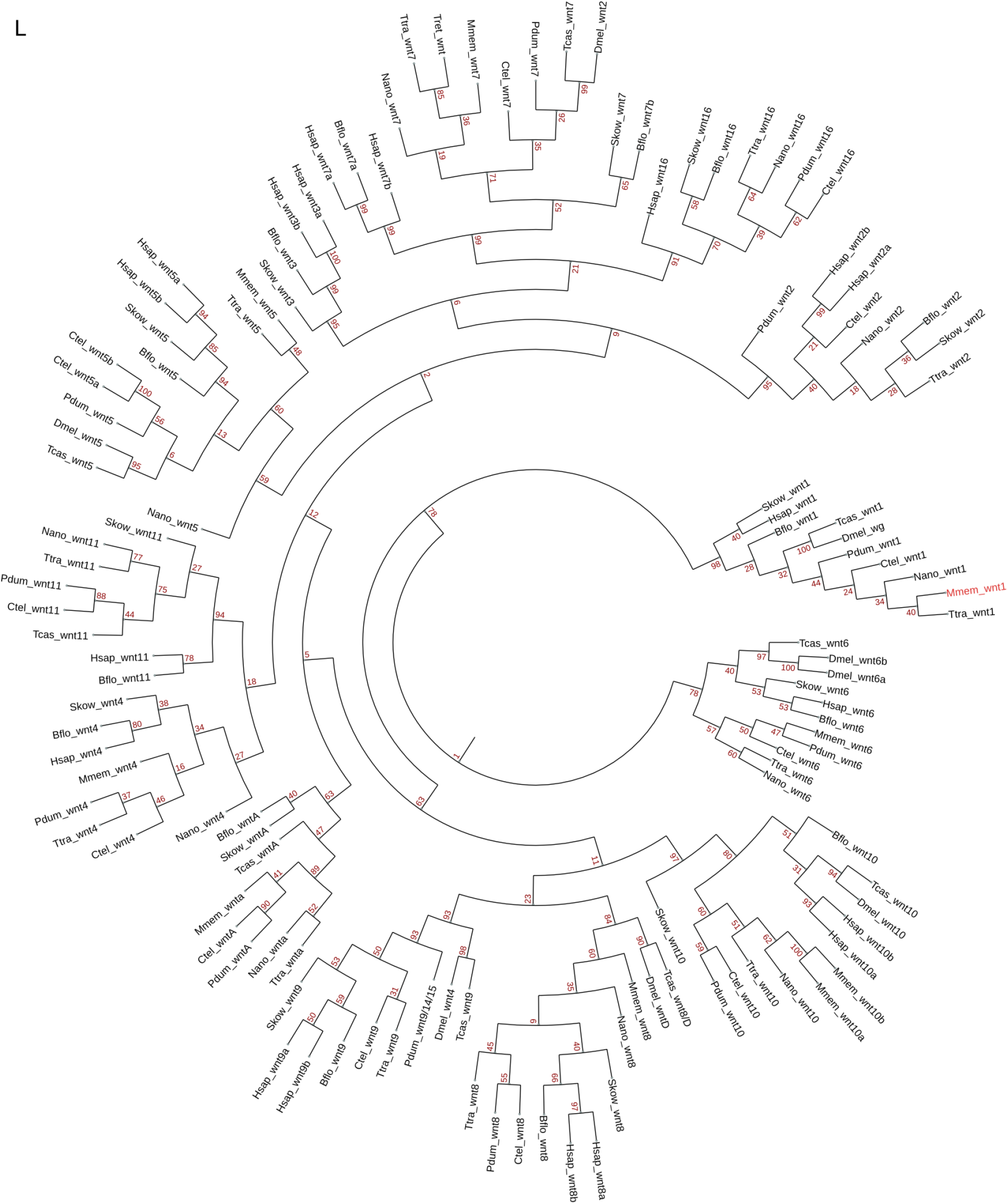
Orthology assignment for the bryozoan genes used in this study. (A) *six3/6*, (B) *dlx* and *evx*, (C) *otx* and *gsc*, (D) *pax6*, (E) *nk2.1*, (F) *foxa*, *foxc* and *foxf*, (G) *nanos*, (H) *bra*, (I) *cdx, twist*, (K) *456*, (L) *wnt 1*. Cladograms show branch support values and bryozoan orthologs in red.

